# A live tumor fragment platform to assess immunotherapy response in core needle biopsies while addressing challenges of tumor heterogeneity

**DOI:** 10.1101/2025.07.18.663728

**Authors:** T.S. Ramasubramanian, Pichet Adstamongkonkul, Christina M. Scribano, Christin Johnson, Sean Caenepeel, Laura C.F. Hrycyniak, Lindsey Vedder, Nicholas Dana, Christian Baltes, Torey Browning, Yuan-I Chen, Thomas Dietz, Evan Flietner, Nicholas Kaplewski, Anna Kellner, Michael Korrer, Chao Liu, Nathan Marhefke, Payton McDonnell, Amreen Nasreen, Victoria Pope, Abhijeet Prasad, Jordyn Richardson, Sidney Schneider, Mikaela Schultz, Chetan Sood, Aishwarya Sunil, Erika von Euw, Eric Wait, Ellen Wargowski, Pooja Advani, Barbara Broome, Antje Bruckbauer, Andrew Godwin, Nima Kokabi, Robert Martin, Mercedes Robaina, Giuseppe Toia, Joshua Routh, Andreas Friedl, Kevin Eliceiri, Mike Szulczewski, Scott Johnson, Jon Oliner, Jérôme Galon, Christian Capitini, Debabrata Mukhopadhyay, Janis Taube, David Braun, Hinco J. Gierman

**Affiliations:** Elephas Biosciences, Madison, WI; Mayo Clinic, Rochester, MN; Orlando Health, Orlando, FL; University of Tennessee Medical Center, Knoxville, TN; The University of Kansas Cancer Center, Kansas City, KS; University of North Carolina, Chapel Hill, NC; University of Louisville Louisville, KY; Advocate Aurora Research Institute, Milwaukee, WI; University of Wisconsin, Madison, WI; Midwestern University, Glendale, AZ; Institute National De La Santé Et De La Recherche Médicale, Paris, France; John Hopkins University, Baltimore, MD; Yale School of Medicine, New Haven, CT

## Abstract

**Background:** Immune checkpoint inhibitors (ICIs) have revolutionized cancer treatment, providing durable and even curative responses. However, most patients do not respond and current biomarkers (eg, programmed death ligand (PD-L1), mismatch repair deficiency (dMMR)/high microsatellite instability (MSI) and tumor mutational burden) lack predictive accuracy. Ex vivo profiling of patient-derived tumor fragments shows promise as a predictive biomarker but relies on substantial surgical tissue to mitigate intra-specimen heterogeneity. Innovations are needed that address these challenges, particularly where limited tissue is available in core needle biopsies (CNBs).

**Methods:** Live tumor fragments (LTFs) were generated from 59 human tumor resections and 31 CNBs from patients enrolled in observational clinical trials (ClinicalTrials.gov identifiers: NCT05478538, NCT05520099, NCT06349642) to assess cytokine induction following ICI treatment. LTFs were encapsulated in hydrogel and cultured ex vivo for up to 72 hours. A sequential treatment strategy that applies control and treatment within the same well was used with response to ICI or αCD3/αCD28 assessed using a multiplex secretome assay. Viability was assessed using established metabolic assays and dynamic optical coherence microscopy.

**Results:** LTFs maintained viability and retained T cells responsive to stimulation throughout ex vivo culture. Multiplex immunofluorescence and immunohistochemistry showed key components of the tumor microenvironment, including relative proportions of CD4+ and CD8+ immune cell populations, were preserved. Specimens positive for PD-L1 or dMMR/MSI-high were enriched for cytokine upregulation, including T-cell response cytokines IFNγ and CXCL10, after αPD-1 treatment. To demonstrate clinical applicability of the sequential treatment strategy, CNBs from patients with lung, gastrointestinal or kidney cancer were profiled and differential cytokine induction in response to ICI treatment was observed.

**Conclusions:** The novel ex vivo platform presented is capable of detecting T-cell response to ICI treatment by using a sequential treatment strategy. This approach addresses challenges associated with cross-well heterogeneity in tissue composition and requires half as much tissue as a cross-well comparison, mitigating tissue limitations typically associated with non-surgical biopsies. Importantly, the platform is compatible with established functional assays as well as non-destructive spatial imaging, enabling researchers to characterize response to ICI longitudinally. Ongoing trials will enable clinicians to assess platform performance in predicting response to immunotherapy.

## BACKGROUND

Immune checkpoint inhibitors (ICIs) have revolutionized cancer therapy, yielding unprecedented clinical benefits for a subset of patients, including curative responses for patients with several types of advanced/metastatic tumors^1^. Currently, 3 clinically validated biomarkers are FDA-approved to predict patient response to ICI treatment: PD-L1 expression^2–6^, microsatellite instability (MSI) / mismatch repair deficiency (dMMR)^5,7^, and tumor mutational burden^8^. Nearly 50% of patients with cancer in the US are not ICI-eligible based on biomarkers^9^. Moreover, of those receiving treatment, only 20% to 30% obtain an objective response^9,10^, underscoring the limited predictive ability of available biomarkers. While anti-programmed cell death 1 (αPD-1) treatment is generally well tolerated, 10% of patients experience severe immune-related adverse events, including fatalities in 0.6% of patients^11,12^. Thus, accurate identification of patients likely to benefit from ICI therapy is critical to optimize clinical outcomes and minimize harm, highlighting the need for more robust, reliable and functionally informative approaches to identify ICI responders.

Ex vivo systems that use live tissues while preserving the native architecture and tumor microenvironment (TME) offer powerful platforms for preclinical studies^13–15^ and have shown promise in predicting therapeutic response^16–18^. Jenkins et al demonstrated cytokine response to PD-1 and CTLA-4 inhibition in patient-derived organotypic tumor spheroids that maintained TME components, such as tumor stroma and immune cells^13^. A key challenge in ex vivo ICI studies is the inherent intra-tumor heterogeneity which leads to unequal tissue composition during tissue allocation for cross-well comparisons. Density and viability of immune and tumor cells can vary drastically across a tumor, resulting in significant variations in cellular composition and microenvironmental conditions between samples of the same tumor specimen^19–21^. This can be problematic with the limited tissue available from a single core needle biopsy (CNB), which is the standard of care clinical diagnostic procedure for advanced-stage metastatic patients who are not surgical candidates and are commonly treated with immunotherapy. Therefore, mitigating intra-specimen variability in tissue composition is essential for reliable ex vivo ICI response assessment.

The optimal platform for assessing ICI response would use live tissue with contiguous TME to enable spatial analysis – such as longitudinal, label-free advanced imaging – and functional response to treatment while eliminating the confounding factor of intra-specimen variability from the limited tissue amounts typical of standard practice diagnostic methods. Here, a novel ex vivo platform was developed for reliable ICI response analysis using live tumor fragments (LTFs) designed to balance the preservation of a functional TME and mitigation of intra-specimen variability.

## METHODS

Methods are summarized below; see detailed methodology in the online supplementary methods.

### Specimens

#### Mouse Models

Experiments using murine models were performed in accordance with the Institutional Animal Care and Use Committee approved protocol (23-003) in an Association for Assessment and Accreditation of Laboratory Animal Care International accredited laboratory per The Guide for Care and Use of Laboratory Animals Eighth Edition^22^. Tumors were removed through a terminal surgical procedure at a volume of 300-500 mm^3^.

#### A. Syngeneic CT26

5×10^5^ CT26 cells (ATCC, CRL-2638) were injected subcutaneously into one or both flanks of female BALB/c mice (The Jackson Laboratory) between 7-8 weeks of age. Cells were tested free of mycoplasma and other pathogens by PCR panel testing (IMPACT II, IDEXX Bioanalytics, 41-00031).

#### B. Humanized PDX

Humanized non-small cell lung cancer patient-derived xenograft (PDX) tumors were generated by warm-passaging 300 µm x 300 µm x 300 µm PDX LTFs into the hind flank of female NSG mice (The Jackson Laboratory) between 7-9 weeks-old. When tumors reached ∼50 mm^3^, mice were injected intravenously with 10-12 x 10^6^ human peripheral blood mononuclear cells from a consenting healthy donor.

#### Human tissue

Protocols for the collection of human specimens were approved by an institutional review board. Resected tumors were collected by a waiver of consent or informed consent and cold temperature shipped overnight. CNBs were collected under informed patient consent from ongoing clinical trials (ClinicalTrials.gov identifiers: NCT05478538, NCT05520099, NCT06349642) and shipped overnight in NanoCool shipping containers (Peli Biothermal, 2-85225). Data from the shipment of 118 consecutive CNBs showed >99% of specimens sustained a temperature between-2°C and 10°C for >98% of the shipping time (Figure S7). For studies including biomarker status, patients were enrolled with colorectal, uterine, head and neck or lung tumors that have FDA-approved indications for αPD-1 treatment based on companion diagnostic (CDx) biomarkers. At the end of the experiment, LTFs were formalin-fixed, and PD-L1 or MMR protein expression immunohistochemical assessment was performed when PD-L1/MMR/MSI status was not available from patient charts.

### Specimen preparation

#### Tumor Resections

Resected tumors were cut into desired dimensions of resection LTFs (eg length x width x thickness of 300 µm x 300 µm x 300 µm) by a fully automated, proprietary cutting device. To prevent T-cell egress^16^, 300 µL of a hydrogel (VitroGel^®^-3, TheWell Bioscience) was added to each well of a 24-well plate and mixed with the LTFs by pipetting. Following polymerization, 1 mL of culture media containing the indicated treatment was added to each well.

#### CNBs

CNBs from 12-to 20-gauge needles were cut using an automated, proprietary cutting device into slices with a thickness of 300 µm. For cross-well comparisons, CNBs were cut perpendicularly (90°) to the longitudinal axis and plated in an 8-well plate, while those for sequential treatment studies were cut at a 20° angle and plated in a 24-well culture plate. Hydrogel (VitroGel^®^-3, TheWell Bioscience; or a proprietary formula, Elephas) was added to each well. After hydrogel polymerization, culture media containing the indicated treatment was added to each well.

### Treatments

ImmunoCult^TM^ Human CD3/CD28 T-Cell Activator (STEMCELL Technologies Inc, 10971) was used at a final concentration of 25 µL/mL, except in Figure 5 (see Supplementary Methods). Immunoglobulin G (IgG) isotype controls and ICI antibodies were used at a concentration of 50 µg/mL (BioXCell). ICI was matched with manufacturer’s recommended IgG control, except in 4 specimens (see Supplementary Methods).

### Assays

#### Viability and cytotoxicity assays

Relative viability was assessed using Cell Counting Kit-8 (CCK8) (Abcam, ab228554). The CCK8 reagent was added immediately after plating, and media was sampled approximately 24 hours later to measure absorbance at 450 nm. Media and reagent were refreshed daily, and the rate of absorbance over time (abs/hour) was calculated. Relative cytotoxicity was measured by the LDH-Glo^TM^ Cytotoxicity Assay kit (Promega, J2381) per manufacturer’s instructions.

#### Optical Coherence Microscopy imaging

A custom-designed optical coherence microscopy (OCM) system was used as previously described^23^ and fitted with a 15X objective lens (Thorlabs, 0.70 NA). Images were acquired at 0, 24, and 48 hours of culture.

#### Cytokine profiling

Conditioned media collected from individual culture wells at defined time points were assessed using the Human XL Cytokine Luminex Performance Assay Premixed Kit per manufacturer’s instructions (R&D Systems, FCSTM18B-30, see Table S4). If the concentration was above the limit of quantitation, the value was set to the upper limit of quantitation. Values below the limit of quantitation were unchanged and removed from analysis where noted. Additionally, six analytes (CD40L, EGF, FGF basic, IL-12p70, IL-15, and FLT-3L) were excluded from analysis due to consistent concentrations below the lower limit of quantitation (LLOQ).

#### Histology

LTFs were fixed in 10% phosphate buffered formalin (Fisher Scientific SF100-4). Resection LTFs were retrieved from the VitroGel^®^-3 post-fixation and transferred to 70% alcohol before being pre-embedded in a 2% agarose solution (Sigma Aldrich, A0576). CNB LTFs encapsulated in Elephas hydrogel were directly transferred into 70% alcohol post-fixation. Resection and CNB LTFs were paraffin-embedded and sectioned at 5-µm thickness on a rotary microtome (Leica, RM2255) and stained with hematoxylin and eosin (H&E) or processed for immunohistochemistry (IHC) or multiplex immunofluorescence (mIF) using Ventana Medical Systems Discovery Ultra Autostainer (see Table S5 for antibodies used).

#### Statistical analysis

Data were reported as mean ± standard deviation and statistical significance assessed by Student’s t-test or Analysis of Variance using Prism 10 software (GraphPad) unless otherwise noted. Fisher’s exact test was performed in the Python SciPy software package^24^. All quantification of histological slides was performed using Visiopharm software.

For CNBs following the sequential treatment protocol, ICI-induced change in cytokine concentrations was determined by the maximum fold change of slope between ICI and IgG control treatment phases (ICI slope/IgG slope).

For resections, ICI-induced change in cytokine concentrations was determined by the difference in cumulative concentrations between ICI and IgG-treated wells. The median absolute deviation (MAD) values were calculated and any sample within two MAD values of the median were then recomputed to generate a trimmed statistic which was transformed into modified Z-scores (Mod. Z) ^25^.

Agglomerative hierarchical clustering was performed to cluster by specimen and analyte using Ward’s method in TIBCO Spotfire™ software. Modified Z-scores of-10 and 10 were selected as the lower and upper saturation points, respectively. A conservative criterion to identify where upregulation occurred was defined by a modified Z-score ≥ 5.0.

## RESULTS

### Ex vivo platform developed to characterize response to ICIs while addressing challenges associated with tumor heterogeneity in live tumor resections

An ex vivo platform was developed in which live tumor resections are cut into cuboid LTFs using an automated proprietary instrument, pooled randomly and encapsulated in a 3D hydrogel (Figure 1A). To optimize fragment size and address intra-tumor heterogeneity, in silico simulations of a CD3-stained melanoma cross-section showed a significant reduction in the coefficient of variation (CV) of CD3+ T-cell numbers in LTFs with edge lengths of 300 (*P <* 0.05) and 100 μm (*P <* 0.01) compared to slices (Figure 1B-C). Although the lowest CV was observed for the 100 µm edge length, 300 µm LTFs were selected for further characterization given the desire to retain as much contiguous TME as possible without compromising on nutrient diffusion^26^. Therefore, 300 μm cuboid LTFs were generated using the automated cutting instrument producing the expected theoretical surface area of 0.09 mm^2^ (theoretical volume of 0.027 mm^3^) across multiple tumor types (Figure S1A-D). These data suggest appropriately controlled cross-well comparisons can be made with pooled LTFs from tumor resections.

**Figure 1.**
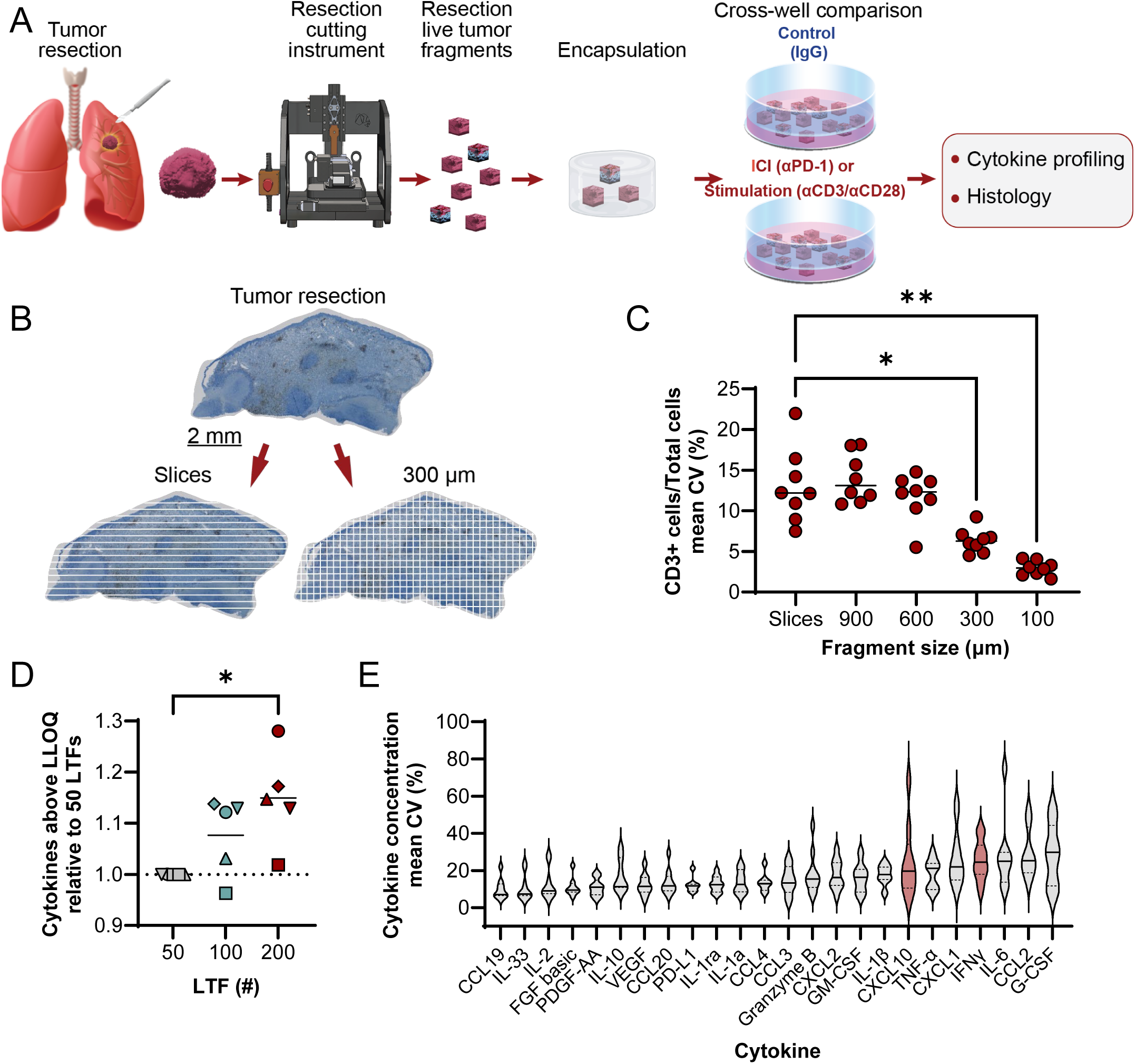
An ex vivo platform for characterizing response to Immune Checkpoint Inhibitors (ICIs) addresses challenges associated with tumor heterogeneity by pooling live tumor fragments (LTFs) created through fragmentation of resections. **A** Schematic shows the workflow for creating LTFs from human tumor resections used for assessing the ex vivo response to treatment. Tumor resections were fragmented using a specialized cutting instrument to create LTFs, encapsulated in a hydrogel and treated with either an ICI (eg, αPD-1), a positive control for T-cell stimulation (eg, αCD3/αCD28), or IgG control, followed by cytokine profiling to assess response to treatment. **B** IHC image of a CD3 stained human melanoma resection showing a standard slice sampling technique (left) compared to LTF 300 µm x 300 µm segments (right). **C** In silico randomized pools of large slices from the melanoma resection showing significantly greater variability in CD3+ staining when compared to randomized pools of LTFs cut to 300 µm edge lengths. **D** Effect of LTF number on the percent of cytokines in the assay panel with measured concentrations exceeding the lower limit of quantitation (LLOQ) following stimulation with αCD3/αCD28 for 24 hours. Data are reported for 5 human tumor specimens (ovarian, n=1; lung, n=1; colorectal, n=1; head and neck, n=2). **E** Variability (CV (%)) across 5 replicate wells of 200 resection LTFs (300 µm (length) x 300 µm (width) x 300 µm (depth)) from each of 10 human tumors (pancreatic, n=2; head and neck, n=4; ovarian, n=3; triple negative breast, n=1) stimulated with αCD3/αCD28. Cytokines of particular interest are CXCL10 and IFNγ (highlighted in red) due to their increase in secretion with T-cell stimulation. * *P* < 0.05, ** *P* < 0.005

To evaluate the effect of LTF number on the variability in T-cell response, LTFs from 5 human tumor resections were dispensed at ∼50, ∼100 and ∼200 LTFs/well. Cytokine induction was assessed following 24-hour stimulation with αCD3/αCD28 (positive control for T-cell response). A reduction in variability was observed across most cytokines with increasing LTF number (Figure S2A-B). Additionally, there was a progressive increase in the number of cytokines exceeding LLOQ as LTF count was increased (Figure 1D). To assess variability in cytokine profiling when using 200 LTFs/well, cytokine concentrations were measured from LTFs derived from 10 human tumor resections following αCD3/αCD28 stimulation (5 replicate wells per tumor). The median CV across all cytokines for each of the individual tumors was below 25% (Figure S2C). A range of variance was observed across individual cytokines, with most cytokines exhibiting an average CV <20% (Figure 1E). This level of variance indicates ∼200 LTFs/well enables the detection of >2-fold changes in cytokine production with statistical significance when paired with multiple replicates.

### Tumor microenvironment, tissue viability and T-cell function of resection LTFs are preserved over 48 hours of ex vivo culture

Cross-section analysis of resected tumors and corresponding LTFs stained with H&E revealed conservation of histological features over 48 hours of culture, demonstrated by the presence of tumor, lymphocytes, fibroblasts, and collagen fibers in both the intact resection specimen (Figure 2A) and the resection LTFs immediately after fragmentation (Figure 2B). Resection LTFs encapsulated and cultured for 48 hours ex vivo showed similar histological features to resection LTFs fixed immediately after cutting (Figure 2C). IHC staining of resection LTFs from 8 tumors representing 5 different tumor types showed reduced total cells (*P*<0.05, Figure S3A-B), reduced CD4+ cells (*P* < 0.05, Figure S3C) and a trend toward reduced CD8+ cells (*P* = 0.07, Figure S3D) after 48 hours of culture. However, the CD4+/CD8+ cell ratio was maintained in all tumor specimens (Figure 2F) with the percentage of CD4+ cells maintained in 5/8 tumors (*P* > 0.05, Figure 2D and S3E) and the percentage of CD8+ cells maintained in 7/8 tumors (*P* > 0.05, Figure 2E and S3F). Taken together, these data suggest the tumor immune microenvironment – particularly the ratio of CD4/CD8 cells – is preserved during ex vivo culture of resection LTFs.

**Figure 2.**
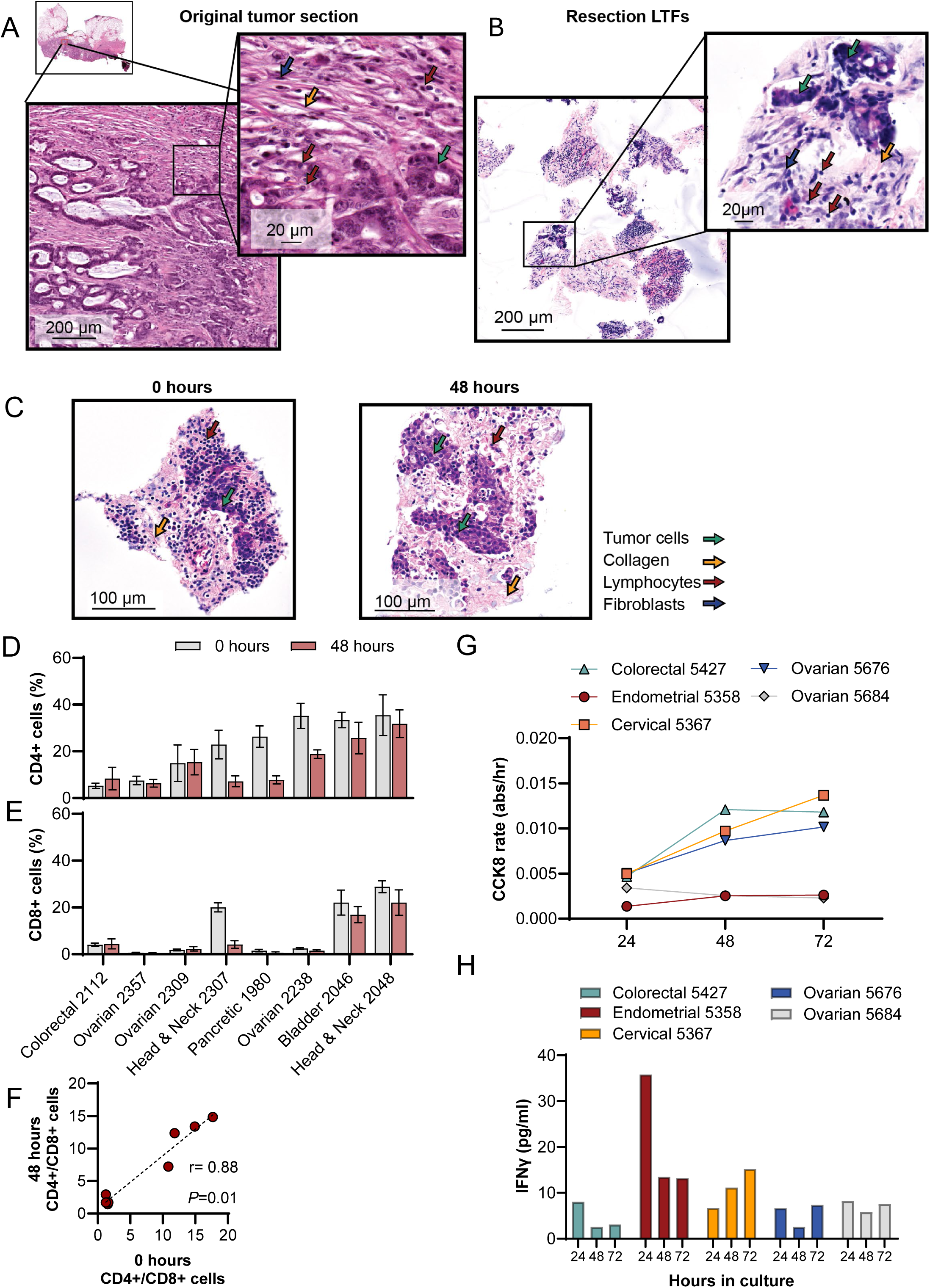
Histological features, tissue viability, the ratio of CD4+/CD8+ cells and cytokines released by activated T cells are preserved in LTFs over 48 hours of ex vivo culture. A, **B** H&E-stained sections of a colorectal cancer resection showing low grade adenocarcinoma. Lymphocytes (red arrows), fibroblasts (blue arrow), tumor cells (green arrow) and collagen fibers (yellow arrow) are present in both the intact resection specimen **(A)** and the resection LTFs at 4 hours **(B).** Comparison of H&E images by a board-certified pathologist identified conservation of histology including intratumor heterogeneity. **C** H&E-stained sections of head and neck LTFs at 4 hours and 48 hours of ex vivo culture showing tumor cells (green arrow), collagen fibers (yellow arrow) and lymphocytes (red arrow). **D-F** Comparison of the percentages of CD4+ (**D**), CD8+ (**E**), and CD4+ to CD8+ cell ratios (**F**) at 4 hours vs. 48 hours as assessed by IHC (N=9 human tumors; colorectal, n=1; bladder, n=1; head and neck, n=2; ovarian, n=3; pancreatic, n=1; uterine, n=1). **G** Changes in viability (CCK8 assay) over 72 hours in culture of LTFs derived from 5 human tumor resections. **H** Changes in T cell function over 72 hours of culture of LTFs as assessed by IFNγ production following 24 hours of stimulation with αCD3/αCD28. * *P* < 0.05.

To further assess the preservation of resection LTFs ex vivo, viability, cell death, and T-cell function were measured in 5 human tumors over 72 hours of culture. Levels of nicotinamide adenine dinucleotide phosphate (NAD(P)H), measured using CCK8^27^, served as a proxy for viability^28^ while lactate dehydrogenase (LDH) release indicated cell death. Figures 2G and S3G show the reciprocal relationship between metabolic activity and cell death over 72 hours in culture. In the first 24 hours, low metabolic rates and relatively high LDH likely reflected damage from tissue processing and/or acclimation to culture conditions. Subsequent time points showed steady or slight increases in metabolic activity and corresponding steady or slight decrease in cell death. This trend continued through 72 hours, demonstrating the culture conditions support cell viability. To assess whether T cells were stimulable, and therefore functional, resection LTFs were stimulated with αCD3/αCD28 at either 0, 24, or 48 hours, and then assayed for interferon gamma (IFNγ) production 24 hours later. Similar IFNγ levels were observed when resection LTFs were stimulated 24 or 48 hours from the start of culture, suggesting T cells are viable ex vivo for at least 72 hours (Figure 2H). Reduced IFNγ production in LTFs stimulated at 24 hours compared to those stimulated at 0 hour is consistent with the decrease in the CD4+ and CD8+ cell populations seen histologically in some samples (Figures S3C-F), as has been observed by others^29^.

### Unsupervised hierarchical clustering groups PD-L1/MMR/MSI biomarker-positive samples amongst those with the greatest cytokine upregulation following ICI treatment

To demonstrate cutting and randomized pooling of LTFs sufficiently addresses intra-tumor heterogeneity and therefore allows for a properly controlled comparison between control and αPD-1 treatments, response to αPD-1 was assessed using LTFs generated from 59 human resections (Table S1, Figure S4A). Clustering of normalized differences in cytokine concentrations (αPD-1 - IgG) revealed groups of tumors with little to no cytokine upregulation (lower Z-scores) in response to ICI on the left and groups of tumors with upregulated cytokines in response to ICI on the right (higher Z-scores) (Figure 3A and S4B). This result suggests increased cytokine production in response to ICI treatment is observed in a subset of tumor specimens.

**Figure 3.**
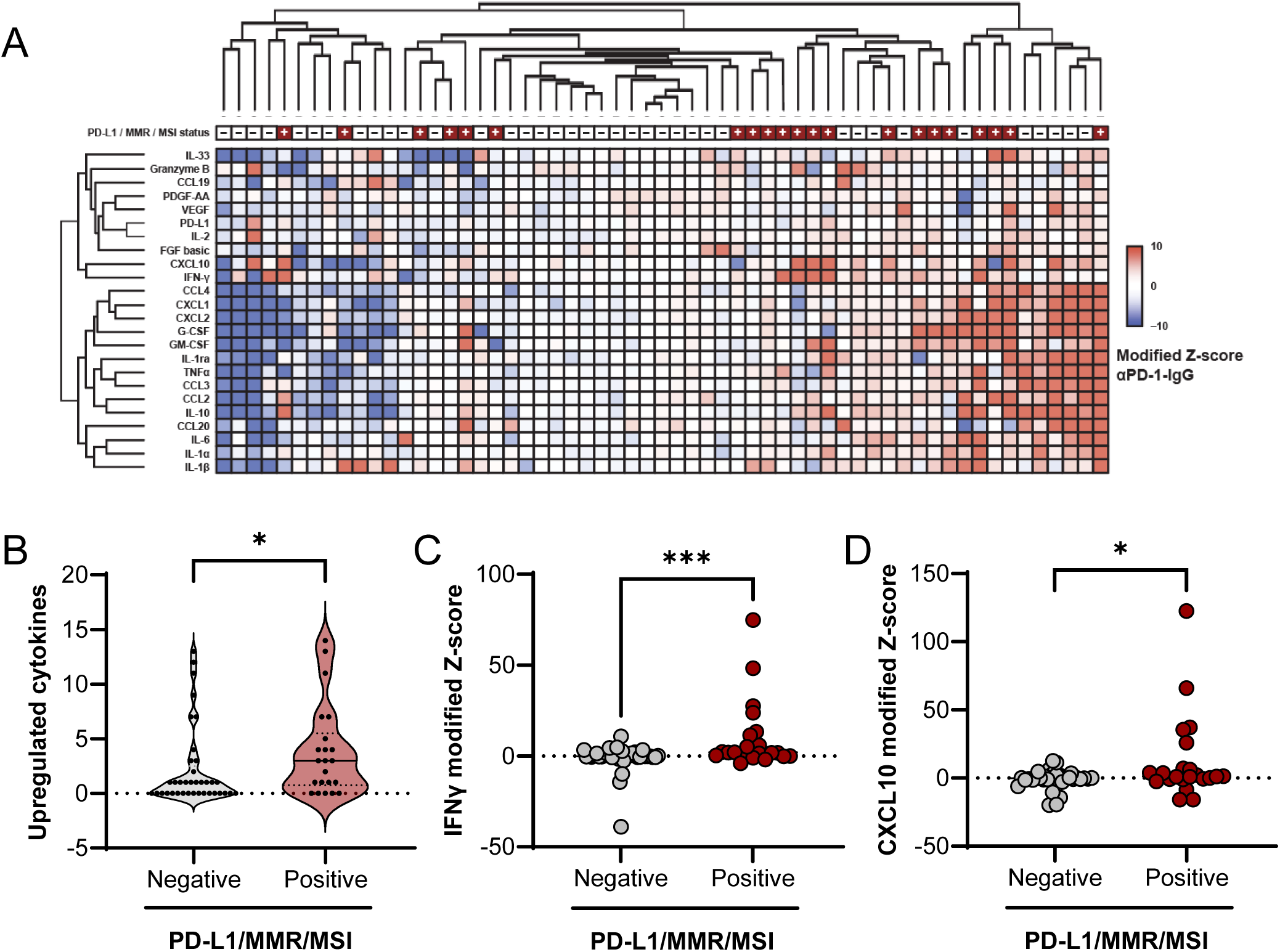
Unsupervised hierarchical clustering of cytokine data groups PD-L1/MMR/MSI-positive specimens amongst those with the greatest cytokine upregulation following ICI treatment. **A** Unsupervised hierarchical clustering of cytokine profiles from patient tumor resections using modified Z-scores of the difference in cytokine concentrations between the ICI and IgG treated groups. To account for variability across wells, replicate wells for each treatment group were run for those specimens where sufficient tissue was available (n=3 for 48 specimens, n=2 for 6 specimens and n=1 for 5 specimens). Positive modified Z-scores (capped at 10) are depicted in shades of red and negative modified Z-scores (capped at-10) are depicted in shades of blue. Specimens with Z-scores of 0 are depicted in white. PD-L1/MMR/MSI-positive specimens are annotated in gray at the top of the heatmap; n=59 specimens. **B** The number of upregulated cytokines for individual specimens, defined by a modified Z-score ≥ 5, is significantly higher in the PD-L1/MMR/MSI-positive cohort compared to the PD-L1/MMR/MSI-negative cohort. Two cytokines known to be associated with T-cell function, IFNγ (**C**) and CXCL10 (**D**), are significantly increased in PD-L1/MMR/MSI-positive compared to PD-L1/MMR/MSI-negative cohorts.

Next, the relationship between ICI-induced cytokine changes and CDx biomarker status (eg, expression of PD-L1 or MMR proteins, or levels of MSI) was evaluated. An enrichment in cytokine upregulation in the PD-L1/MMR/MSI-positive (PD-L1+/dMMR/MSI-high) population compared to the PD-L1/MMR/MSI-negative (PD-L1-/pMMR/MSS) population was observed (Figure 3B). Further analysis of individual T-cell response cytokines showed significant enrichment of IFNγ and C-X-C motif chemokine ligand 10 (CXCL10) upregulation (higher Z-scores) in the biomarker positive samples compared to the biomarker negative samples (Figures 3C-D and Table S2). These data suggest automated tissue fragmentation and randomized pooling of resection LTFs address tumor heterogeneity and may allow one to effectively model ICI treatment response in human tumor resections.

### CNBs with limited tissue availability are less amenable to fragment-pooling approach to overcome tissue heterogeneity

Multiple replicates of control and treated wells enable meaningful cross-well comparisons when sufficient tissue is available, addressing heterogeneity in tumor resections. However, CNBs, used in standard practice during clinical diagnosis, pose a challenge due to scarcity of tissue available to address intra-tumor heterogeneity, making conventional cross-well comparison requiring multiple replicates less feasible. To confirm this, we compared control (IgG) and treated (αPD-1) cytokine concentrations between pooled LTFs from 59 human tumor resections with pooled LTFs from 24 CNB specimens in a cross-well manner. To assay multiple wells from a single CNB specimen, CNBs were cut at a 90° angle to maximize the number of CNB LTFs obtained (Table S3). As a result, 3–5 CNB LTFs were plated per well, with triplicate wells prepared for each condition. The media amount per well was reduced to maintain a consistent tissue-to-media volume ratio to that of experiments using resection LTFs. While the number of quantifiable cytokines in the tumor resections was similar to those from the CNB specimens for IgG controls (Figure 4B), there was significantly greater variation in ICI-induced cytokine production in the LTFs from CNB tumors compared to resection tumors (Figure 4C, Figure S5), suggesting heterogeneity was not fully addressed in pooled LTFs from CNBs. The limited amount of tissue from a single CNB (eg, an 18 gauge of ∼10 mm in length) is roughly equivalent to the tissue amount used in a single well of 200 resection LTFs (total tissue volume about 5.4 mm^3^) and cannot support the 6 replicate wells that would be needed for a meaningful cross-well comparison (Figure 3). These results suggest intra-specimen heterogeneity is not mitigated in pooled CNB LTFs, and tissue scarcity does not allow for a robust cross-well comparison between control and treatment regimens.

**Figure 4.**
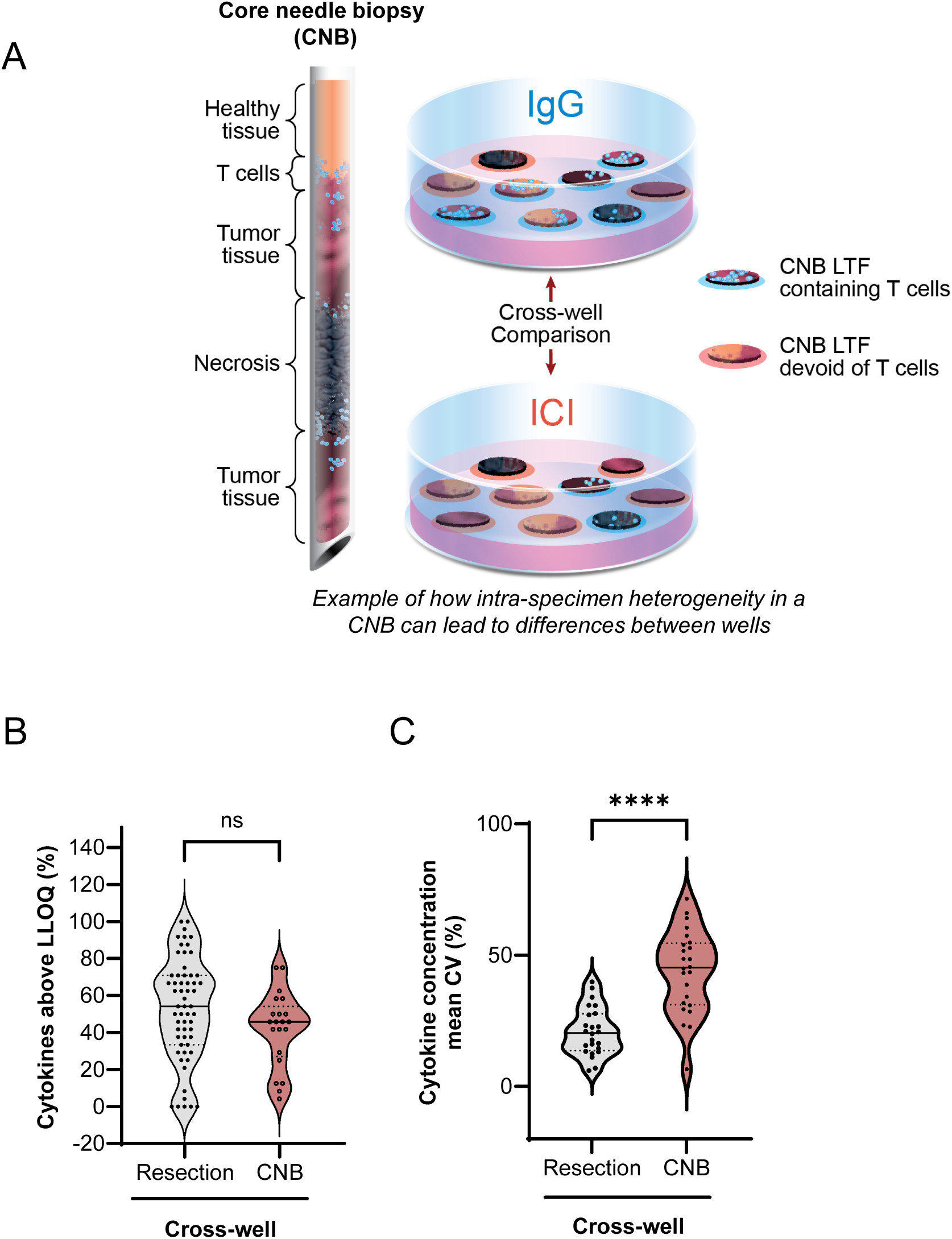
**Cross-well comparison of cytokine production in response to ICI treatment is not amenable to CNBs due to limited tissue availability and inability to address intra-specimen heterogeneity**. **A** Schematic showing the distribution of tissue from a core needle biopsy (CNB) in an experiment using the cross-well configuration where tissue amount is insufficient to address intra-specimen heterogeneity. **B** The percentage of cytokines out of a 30-cytokine panel with concentrations above LLOQ for the cross-well configuration for resection LTFs (Resection cross-well) and CNB LTFs (CNB cross-well) following treatment with IgG control for 24 hours. There is no significant difference in the ability to detect cytokines between the two form factor protocols (ns = not significant, *P* = 0.089). **C** The variability in concentrations of cytokines from resection LTFs and CNB LTFs plated in the cross-well configuration and treated with ICI for 24 hours. There is significantly greater variability in cytokine concentrations between replicate wells of treated groups in CNB LTFs compared to resection LTFs. **** *P* < 0.0001

**Figure 5.**
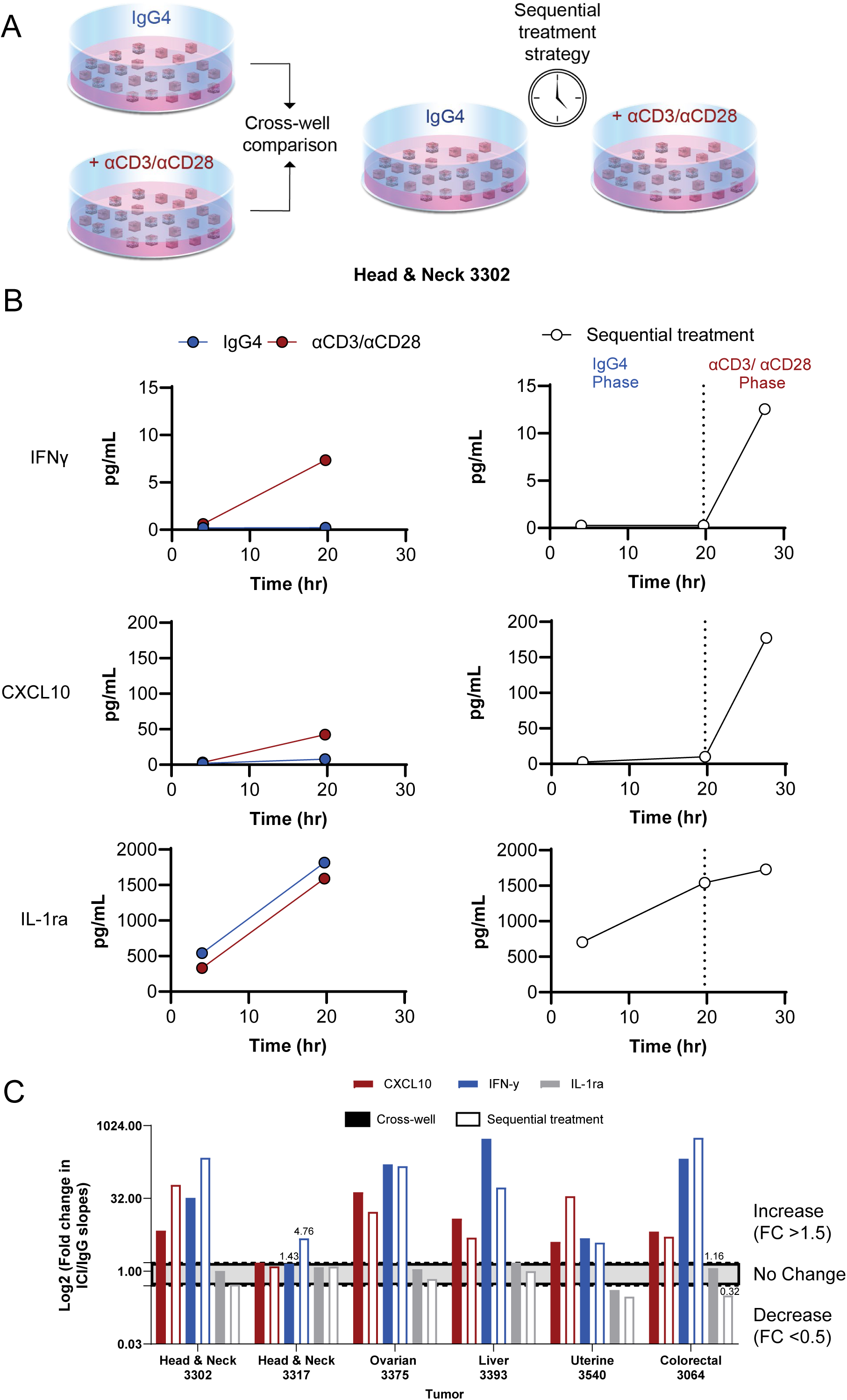
Challenges associated with intra-specimen heterogeneity can be addressed using a sequential treatment strategy where changes in cytokine production rate are compared between treatment and control in the same tissue. **A** Schematic of protocol used to validate sequential treatment strategy. Tumor resections were used due to the availability of sufficient tissue to address inter-well heterogeneity in tissue composition and enable meaningful cross-well comparisons. Tumor resections were cut into resection LTFs, encapsulated in hydrogel and treated with IgG for 28 hours (shown in blue), positive control αCD3/αCD28 to stimulate T cells for 28 hours (shown in red), or IgG4 for the first 20 hours and αCD3/αCD28 for the final 8 hours of culture (shown in red). Conditioned media was sampled from each of the wells at 4, 20, and 28 hours for cytokine profiling. **B** IFNγ induction in a head and neck human tumors specimen. IFNγ and CXCL10 induction was observed following αCD3/αCD28 stimulation in cross-well (left) and sequential treatment (right). IL-1Ra remained unchanged following αCD3/αCD28 stimulation. **C** To characterize the similarity in cytokine production following the cross-well and sequential treatment experimental designs, the fold change in rates (slopes) of cytokine production following αCD3/αCD28 stimulation were calculated compared to rates of cytokine production during IgG exposure. The fold changes (FC) in rates of cytokine production for cross-well and sequential treatment strategies were compared to determine similarities of magnitude and direction (Increase >1.5; no change = 0.5-1.5; decrease <0.5). IFNγ, CXCL10 and IL-1Ra showed 100%, 83% and 83% concordance, respectively, between cross-well and sequential treatment strategies. Numerical labels are shown in cases when cross-well and sequential treatment data are discordant. Grey band indicates no change.

### Challenges associated with tumor heterogeneity in CNBs can be addressed using a sequential treatment strategy

To address tumor heterogeneity and the compounding tissue scarcity of CNBs, a sequential treatment strategy was devised wherein CNB LTFs were treated sequentially with the negative control followed by ICI within the same well. The treatment effect was evaluated via longitudinal assessment of cytokine production rates. Here, the tissue was stimulated after a control-phase in the sequential treatment setting to determine if it elicits a similar response as stimulating immediately after encapsulation. For these experiments, 300 µm resection LTFs at a target of 200 LTFs/well were used to address intra-specimen heterogeneity and enable a meaningful comparison between conditions in a cross-well setting. Resection LTFs were either treated within dedicated wells for 28 hours with IgG or αCD3/αCD28 (cross-well), or within the same well for 20 hours with IgG followed by αCD3/αCD28 for an additional 8 hours (sequential treatment) (Figure 5A). Longitudinal, cumulative concentration assessment of CXCL10 and IFNγ from a representative human head and neck tumor resection showed an increased production rate with αCD3/αCD28 stimulation for both cross-well and sequential treatment experiments (Figure 5B). Inclusion of 5 additional tumors showed a similar trend with 100% concordance between cross-well and sequential treatment with increases in the slopes of IFNγ and CXCL10, in response to αCD3/αCD28 treatment over IgG. While the magnitude of increase in IFNγ and CXCL10 in response to αCD3/αCD28 between cross-well and sequential treatment varied, the directionality of response was in high agreement with 5/6 tumor samples showing >1.5 fold change for IFNγ and all tumor samples showing >1.5 fold change for CXCL10 (Figure 5C). Additionally, interleukin-1 receptor agonist (IL-1Ra), a regulatory cytokine, was well expressed at baseline and did not change significantly following αCD3/αCD28 treatment. For this reason, IL-1Ra is used to highlight a ubiquitously expressed, non-responding cytokine in the analysis.

### Longitudinal imaging is enabled with CNB LTFs when using a specialized cutting instrument and encapsulation in a proprietary hydrogel

Leveraging the advantages of the sequential treatment strategy, which does not require multiple wells, LTF generation was adapted, shifting from creating cuboid LTFs optimized for efficient randomization to producing larger LTFs with a greater area of contiguous tissue architecture that allows for better assessment of spatial tissue organization and cell interactions (Figure 6A).

**Figure 6.**
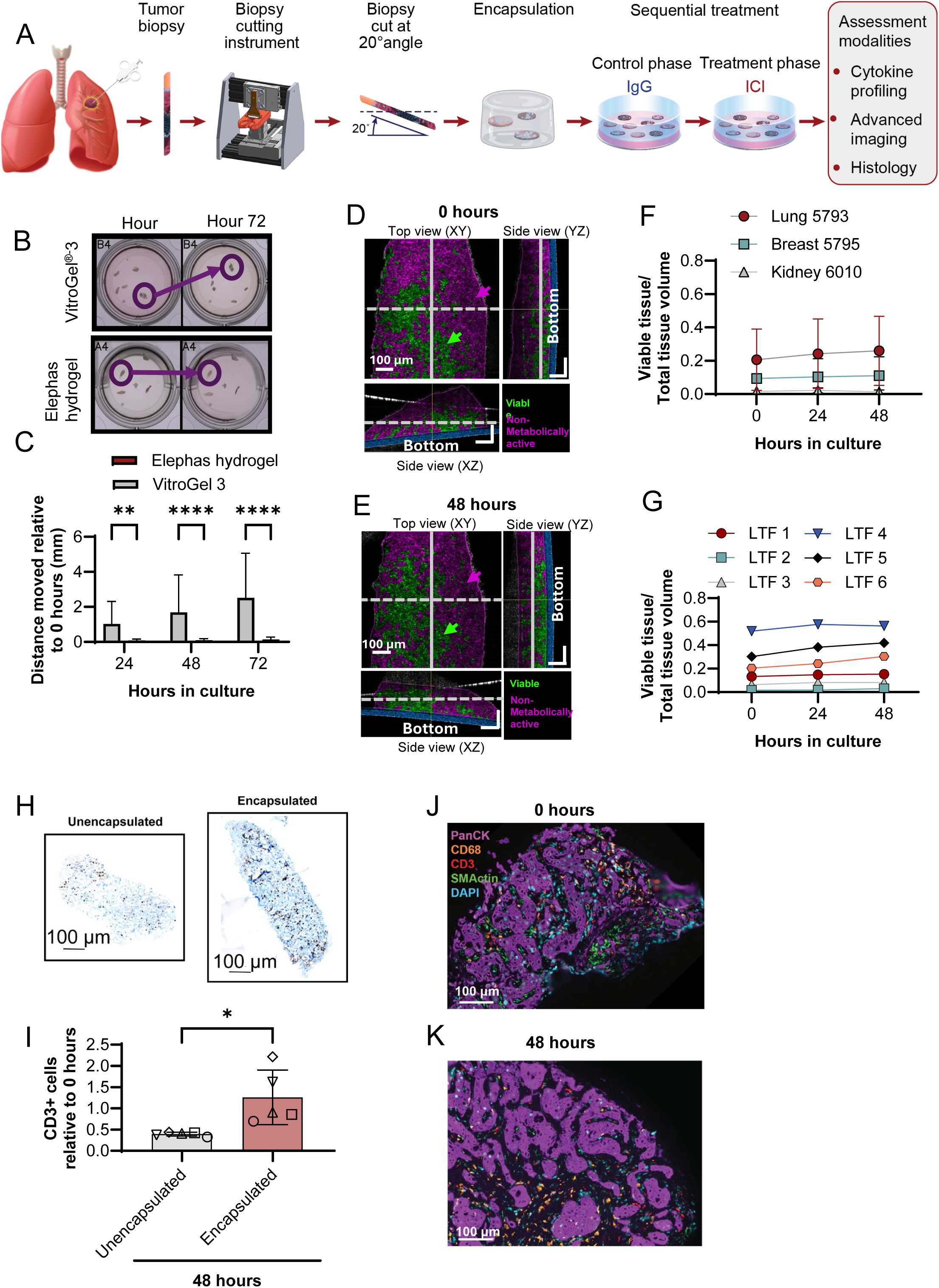
Creation of LTFs from CNBs using a specialized cutting instrument and subsequent encapsulation in a proprietary hydrogel enables longitudinal advanced imaging and retains T cells. **A** Schematic shows the workflow for creating LTFs from human CNBs used for assessing the ex vivo response to treatment. Tumor CNBs are sliced at a 20° angle using a specialized cutting instrument to create LTFs which are then encapsulated in a proprietary hydrogel (referred to as Elephas hydrogel hereafter) for ex vivo culture, enabling cytokine profiling, histological analysis and advanced imaging. **B** The spatial stability of CNB LTFs created from CT26 syngeneic tumors (N=4 tumors) over 48 hours of ex vivo culture when encapsulated in VitroGel^®^-3 versus Elephas hydrogel. Circles highlight regions of hydrogel where LTF stability is compromised with VitroGel-3 encapsulation but not with Elephas hydrogel. **C** The distances between CNB LTF locations at 72 hours vs. 4 hours of culture were measured and reported relative to 4 hours when encapsulated in Elephas hydrogel or VitroGel^®^-3. Limited movement was observed with Elephas hydrogel, highlighting improved stability and enabling longitudinal advanced imaging. **D-E** Dynamic optical coherence microscopy (dOCM) imaging of a human lung (NSCLC) CNB LTF at 4 hours (**D**) and 48 hours (**E**) in culture. Volume positive for metabolic activity indicating tissue viability is false-colored in green and volume negative for metabolic activity is false-colored in magenta. Note that volume negative for metabolic activity represents non-viable and acellular tissue. **F** The ratios of viable voxels to total voxels are reported across 3 human tumors (lung, breast, kidney) and show maintenance of viability over 48 hours in culture. **G** The ratio of viable volume (measured in voxels) to total volume reported for 6 CNB LTFs from Lung 5793 highlights heterogeneity in viability across LTFs from a single CNB. **H** A significant decrease in the number of CD3+ cells, normalized to area, is seen in unencapsulated compared to encapsulated humanized PDX CNB LTFs after 48 hours in culture (each symbol represents 1 tumor, N=5 tumors per group, * *P* < 0.05). **I** Representative CD3-labeled sections of humanized PDX CNB LTFs from encapsulated (top) and unencapsulated (bottom) tissues after 48 hours of ex vivo culture. **J-K** Multiplex immunofluorescence labeled fragments from Liver 5605 show the presence of tumor cells (PanCK), macrophages (CD68), T cells (CD3), stroma/fibroblasts (SMActin) and cell nuclei (DAPI) at 4 hours (**J**) and 48 hours (**K**) in culture.

Assessment of cellular features and interactions in cultured tissue over time by imaging requires the ability to register tissue loci easily for repeated longitudinal assessment. To reduce fragment movement, a proprietary hydrogel (Elephas hydrogel) was designed and tested against a commercially available hydrogel (VitroGel^®^-3). CNB LTFs generated from a CT26 syngeneic mouse tumor were encapsulated in either VitroGel^®^-3 or Elephas hydrogel and cultured for 72 hours (Figure 6B). The positions of LTFs were tracked over time and the magnitude of movement was quantified for the 2 hydrogels. CNB LTFs embedded in the commercial hydrogel showed significant drift over time in culture while CNB LTFs encapsulated in Elephas hydrogel were stable (Figure 6C). dOCM, which enables measurement of intra-cellular activity (eg, mitochondrial movement), was employed to longitudinally evaluate viability through measuring metabolic activity of LTFs from 3 human CNBs over a 48-hour period. Representative dOCM images from 0 hours and 48 hours, where viable/metabolically active tissue (Viable, false-colored in green) and metabolically inactive tissue (Non-metabolically active, false-colored in magenta) were visualized (Figures 6D-6E). Over the course of culture, the tissue-maintained viability with a similar proportion of viable tissue area between 0 and 48 hours (Figure 6F). Quantitative analysis of the 6 individual CNB LTFs from the same non-small cell lung cancer specimen further supports this trend by showing the maintenance of viable tissue over 48 hours in culture (Figure 6G). The variation in viable tissue proportions in these CNB LTFs at hour 0 highlights CNB heterogeneity. These findings support the compatibility of the platform with dOCM as a non-invasive technique to track metabolic changes in viable tissue over time.

### CNB LTFs encapsulated in Elephas hydrogel retain cellular components of the TME over 48 hours of ex vivo culture

To demonstrate that Elephas hydrogel retains T cells within CNB LTFs, IHC staining was performed to identify CD3+ T cells in encapsulated and un-encapsulated CNB LTFs from humanized PDX tumors after 48 hours of ex vivo culture (Figure 6H). The proportion of T cells retained within CNB LTFs after 48 hours in culture relative to 0 hours was significantly higher in encapsulated CNB LTFs than unencapsulated LTFs (*P* < 0.05, Figure 6I).

Similar to resection LTFs, a steady or increasing rate of viability as assessed by CCK8 over 72 hours of culture (Figure S8A) and a concurrent steady or decreasing rate of LDH accumulation following an initial spike within the first 24 hours (Figure S8B) were observed. Assessment of CNB LTFs by H&E staining showed retention of major histological features between 0 and 48 hours in culture (Figure S8C), similar to the preservation in resection LTFs observed earlier (Figure 2C). In addition, mIF confirmed the preservation of cellular composition in LTFs, including T cells, macrophages, tumor cells and fibroblasts (Figures 6J-K). Together, the retention of histological features as well as the results from metabolic and cytotoxicity assays suggest the overall viability of CNB LTFs is maintained throughout ex vivo culture similar to what was observed in cuboid LTFs from resections (Figure 2). Changing the form factor from 300 μm cuboid LTFs to larger elliptical slices did not have an appreciable effect on the ability to culture and maintain the TME.

### The sequential treatment strategy allows for characterization of ICI response ex vivo in CNBs

The efficacy of the sequential treatment strategy was determined in diagnostic CNBs obtained from patients enrolled in three ongoing clinical trials. After CNBs were sliced on a proprietary cutting instrument, LTFs were plated, encapsulated, and cultured for 48 hours to assess cytokine response to ICI. LTFs were treated with IgG at the start of culture and then with αPD-1 (lung specimens), αPD-L1 (gastrointestinal, GI specimens), or αPD-1 + αCTLA-4 (kidney specimens) at around 20 hours, such that LTFs in each well received control and ICI treatment sequentially (Figure 7A). Variable cytokine response profiles across specimens were observed. For example, in Lung 4624, Colon 4834, and Kidney 5023 there was an increase in the rate of accumulation (slope) of both IFNγ and CXCL10 after ICI treatment, relative to IgG control (Figure 7B, 7E and 7H, top and bottom, respectively). Conversely, in Lung 4578, Esophageal 4798, and Kidney 5327 the rates of accumulation (slope) of both IFNγ and CXCL10 remained unchanged following ICI treatment relative to control (IgG) (Figure 7C, 7F and 7I, top and bottom, respectively). In contrast to the robust increases observed with T-cell response cytokines IFNγ (max FC = 12.4) and CXCL10 (max FC = 19.2), IL-1Ra showed a limited change in slopes between the treatment and control phases (max FC = 3.9, Figure 7D, G, J and Figure S8A-D). suggesting specific changes in immune related cytokines can be measured using the sequential treatment approach following treatment with ICI.

**Figure 7.**
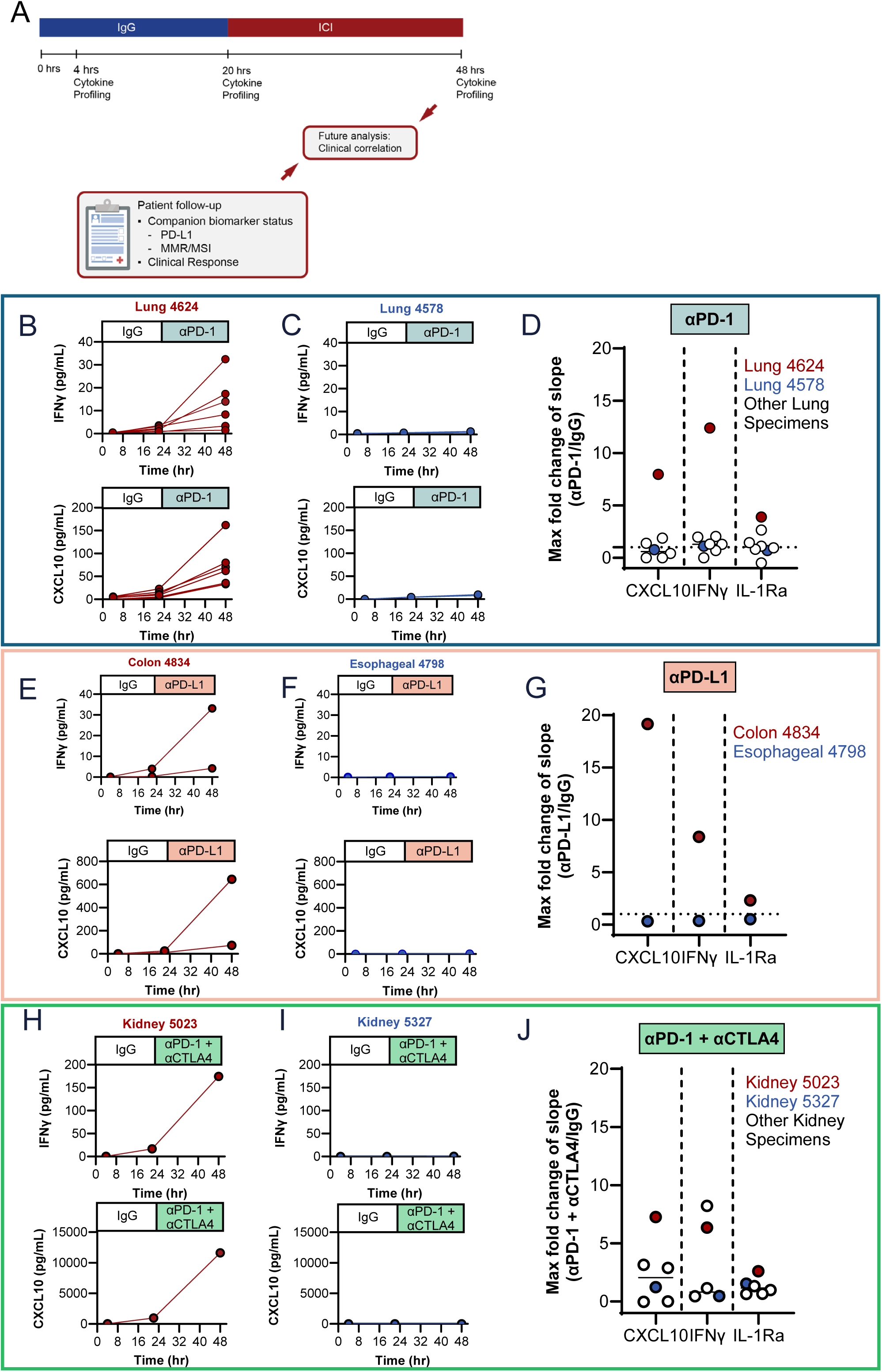
A sequential treatment strategy allows for characterization of ICI response ex vivo in CNBs where tissue is limiting. **A** Encapsulated LTFs from CNBs are sequentially treated with IgG control followed by ICI. Conditioned media is collected at the ∼4-, ∼20-and ∼48-hour time points to enable measurement of cytokine concentrations which are used to calculate the rate of cumulative cytokine production during the control (4-20 hours) and ICI (20-48 hours) treatment phases. Fold change in the rate of cytokine production (slope of ICI treatment phase / slope of IgG control phase) can be used to assess ex vivo treatment response. Exemplary human lung (**B**, Lung 4624 treated with αPD-1), colon (**E**, Colon 4834 treated with αPD-L1) and kidney (**H**, Kidney 5023 treated with αPD-1 + αCTLA-4) cancer CNBs exhibiting an increase in rate of IFNγ and CXCL10 production following ICI treatment suggesting a potential cytokine response. Exemplary human lung (**C**, LU4578 treated with αPD-1), esophageal (**F**, Esophageal 4798 treated with αPD-L1) and kidney (**H**, Kidney 5327 treated with αPD-1 + αCTLA-4) cancer CNBs exhibiting a similar rate of IFNγ and CXCL10 production between the ICI and IgG treatment phases suggesting an unlikely cytokine response. Note that multiple CNBs were obtained from patients 4624, 4578 and 4834 enabling profiling of replicate wells. The fold change in the slope of the ICI treatment phase compared to the slope in the IgG treatment phase for IFNγ and CXCL10 across 7 lung (**D**), 2 gastrointestinal (**G**) and 6 kidney (**J**) cancer CNBs. Individual data points report maximum fold change for those specimens where multiple CNBs/replicate wells were profiled. Note that the maximum fold change for the exemplary cytokine responsive specimens (Lung 4624, Colon 4834 and Kidney 5023, shown in red) is at the top of the distribution while the maximum fold change for the exemplary cytokine nonresponsive specimen (Lung 4578, Esophageal 4798 and Kidney 5327, shown in blue) is close to 1.

## DISCUSSION

The ex vivo platform described here enables characterization of cytokine response to ICIs while addressing the challenges of tumor heterogeneity and limited tissue availability in clinically relevant CNBs. By preserving the native TME, this approach maintains the architectural and cellular context essential for an accurate assessment of immune response. The sequential treatment strategy, wherein CNB LTFs are exposed to a control followed by ICI in the same well, enables paired comparisons within individual replicates. This design mitigates sampling bias, maximizes tissue usage, and facilitates assessment of ICI-induced cytokine induction in a short timeframe, even in biopsies with limited material.

An intact TME is increasingly recognized as being crucial for assessing ICI response ex vivo, due to the complex interactions among its diverse cell types that influence treatment outcomes. Components of the TME, such as cancer-associated fibroblasts^30^, tumor-associated macrophages^31^, T cells^32^ and the tumor cells themselves^33^, have been shown to be intricately involved in orchestrating ICI response. Furthermore, the spatial organization and interactions of these cells within the TME are crucial for effective immune responses and subsequent ICI efficacy^34–36^. Resection and CNB LTFs preserve the native TME, retaining histological features such as fibroblasts, tumor cells, and immune cells—including T cells and macrophages— through the end of the culture period. T cells retained in LTFs encapsulated with the Elephas hydrogel remain responsive to stimulation, allowing for the serial assessment of cytokine induction rates following ICI treatment. More importantly, T cells are exposed to ICI treatment within the context of the native TME in large contiguous sections of tumor tissue, thus preserving the complex spatial relationships that influence treatment efficacy.

Representation of tumor heterogeneity is both critical for accurate prediction of response to immunotherapy and problematic to cross-well experimentation through the introduction of variation resulting from tumor sampling. It is therefore important to represent the heterogeneity of tumors, while reducing the potential for high variance, which can confound the measurement of treatment effects. To address this, the resection-based protocol generates randomized pools of cuboid LTFs to support cross-well cytokine comparisons. However, cytokines known to be induced by immune cell activation, such as IFNγ and CXCL10, show higher variances, likely a result of the differential distribution of T cells and other immune cells across tissue. This necessitates the use of multiple replicates for robust statistical evaluation. The platform’s ability to maintain tumor heterogeneity and the native TME was demonstrated in human tumor resections, where ICI-induced cytokine responses were associated with PD-L1 and MMR/MSI status. A perfect alignment of upregulated cytokine clustering among biomarker-positive tumors was not observed, as expected given the imperfect accuracy of current biomarkers^2–7^. Future studies assessing the correlation between ex vivo and clinical responses will determine whether the platform can better predict response to ICI therapy Preliminary experiments with CNBs from lung, kidney, and gastrointestinal tumors revealed cytokine induction patterns unique to individual specimens, particularly in IFNγ and CXCL10. In cases where CNB tissue amount supported multiple wells for assessment, intra-specimen heterogeneity was evident from the range of responses to ICI observed. However, the sequential treatment approach allows for assessment of response within each individual replicate well such that a positive response in any replicate well suggests potential treatment sensitivity, while the absence of response across all replicates indicates potential resistance.

While this platform offers significant advantages, some challenges warrant mention. First, necrotic biopsy samples pose a risk of misinterpretation, potentially appearing as non-responders and introducing false negatives. Colorimetric assays to assess viability can be integrated into the platform to mitigate this, however they may negatively affect the tissue over time^28^. Adding the viability assay downstream of sequential treatment may be an approach to mitigate the negative effects of these assays, however the effects of treatment on this assay need to be elucidated. A better approach that is dye-free and non-invasive would be the use of dOCM, which detects intracellular movement to assess tissue viability. When integrated at the assay start, dOCM could enable the exclusion of necrotic samples from analysis. Second, the presence of tumor-free samples, another source of potential false negatives, necessitates careful identification. While currently reliant on histological assessment, future advancements in imaging technologies may offer more efficient solutions for excluding samples lacking tumor content. Finally, collecting CNBs from a tumor may not always accurately reflect the potential for the patient’s tumor to respond to ICI due to tumor heterogeneity. In addition, CNBs are not always collected across multiple lesions. Previous studies have demonstrated differential ex vivo responses to ICI from within a tumor as well as across multiple lesions within a patient^16^. Ideally, the entire tumor would be assessed when possible, and it is critical to assess all available biopsy tissue to mitigate tumor heterogeneity. Additionally, a positive control for T-cell presence—such as αCD3/αCD28 stimulation following ICI treatment—could mitigate this risk to some extent, confirming the presence of functional T cells and a non-response.

Other ex vivo models for predicting ICI response preserve the native TME to varying degrees. Air-liquid interface organoids preserve the intratumoral T-cell repertoire and have been shown to upregulate IFNγ, granzyme B and perforin following PD-1 blockade^37^. However, such organoid models often involve expansion of cells in culture, limiting their clinical utility for guiding treatment decisions. Tumor fragment models derived from surgical resections also preserve the native TME and allow cytokine-based assessment of response ^13,16,17^. These models rely on pooling large tissue amounts to mitigate heterogeneity and are not compatible with biopsies, excluding unresectable, advanced and metastatic patients most likely to receive ICI therapy from potential evaluation.

The sequential treatment strategy presented here enables detection of cytokine changes in response to ICI exposure in CNBs. Ongoing clinical studies are targeted to enroll over 600 patients and designed to evaluate the extent of correlation between ex vivo cytokine response and clinical outcomes, validating this approach as a predictive tool (ClinicalTrials.gov identifiers: NCT05478538, NCT05520099, NCT06349642). By leveraging CNBs and a sequential treatment design, the platform supports longitudinal profiling of tumor response and offers a practical method for guiding personalized ICI therapy. Importantly, this approach is not limited to ICI monotherapies. The platform also has the potential to evaluate other immunotherapy modalities, including immune cell–based therapies and combination regimens. These applications could significantly expand the platform’s clinical utility. Further studies are ongoing to explore these opportunities and validate use beyond ICI treatment.

## DECLARATIONS

### Availability of data and material

Data sets used in the current study are available on reasonable request.

## Acknowledgements

This paper is dedicated in the memory of Abhijeet Prasad, who made significant contributions to this work.

We express our sincere gratitude to the patients, clinical staff, and researchers at participating clinical sites, as well as to current and former Elephas employees and scientific collaborators, for their valuable contributions to this work.

Tissue samples were provided by the Cooperative Human Tissue Network which is funded by the National Cancer Institute.

The authors thank the University of Wisconsin Carbone Cancer Center BioBank, supported by P30 CA014520, for use of its facilities and services and Mercy Research located at Mercy Sindelar Cancer Center led by principal investigator Peter DiPasco, MD.

The authors acknowledge the support of The University of Kansas Cancer Center’s Biospecimen Repository Core Facility staff, funded in part by the National Cancer Institute Cancer Center Support Grant P30 CA168524, The University of Louisville, Division of Surgical Oncology Biobank and The University of Tennessee Medical Center’s Biobank for its service in providing biospecimens under the direction of Antje Bruckbauer, MD, PhD.

## LIST OF ABBREVIATIONS

αPD-1: anti-programmed cell death-1
αPD-L1: anti-programmed cell death ligand-1
CCK8: cell counting kit 8
CDx: Companion diagnostic
CD3/CD28: cluster of differentiation 3/28
CNB: core needle biopsy
CTLA4: cytotoxic T-lymphocyte-associated protein 4
CV: coefficient of variation
CXCL10: C-X-C motif chemokine ligand 10
dMMR: deficient DNA mis-match repair
dOCM: dynamic Optical Coherence Microscopy
DPBS: Dulbecco’s phosphate buffered saline
FBS: fetal bovine serum
GI: gastrointestinal
H&E: hematoxylin and eosin
ICI: immune checkpoint inhibitor
IF: immunofluorescence
IFNγ: interferon gamma
IgG: immunoglobulin G
IHC: immunohistochemistry
IL-1Ra: interleukin-1 receptor agonist
LDH: lactate dehydrogenase
LLOQ: lower limit of quantitation
LTF: live tumor fragment
MAD: median absolute deviation
mIF: multiplexed immunofluorescence
MMR: DNA mis-match repair
MSI: microsatellite instability
NAD(P)H: nicotinamide adenine dinucleotide phosphate
NSCLC: non-small cell lung carcinoma
PBMC: peripheral blood mononuclear cell
PDX: patient-derived xenograft
pMMR: proficient
DNA: mis-match repair
TME: tumor microenvironment
WST-8: water-soluble tetrazolium salt

