## Supplemental methods for "A live tumor fragment platform to assess immunotherapy response in core needle biopsies while addressing challenges of tumor heterogeneity"

### Specimens

Mouse Models:

Experiments using murine models were performed in accordance with the Elephas Institutional Animal Care and Use Committee approved protocol (23-003) in an Association for Assessment and Accreditation of Laboratory Animal Care International accredited laboratory per The Guide for Care and Use of Laboratory Animals Eighth Edition [22]. Female mice received at about 6 weeks-old were housed in a 12/12-hour light/dark cycle, provided ad libitum access to food and water and supplied with cage-enrichment devices (eg, nesting tubes, exercise wheels, etc.). During tumor growth, mice were monitored for overall health through daily mass measurements and investigator observation. Tumor dimensions were measured twice weekly using calipers to determine length and width and to estimate volume ($V = 0.5\ L \times W^2$). Tumors were surgically removed at a volume of 300-500 $mm^3$.

Syngeneic CT26:

CT26 mouse colon carcinoma cells (ATCC, CRL-2638) were cultured at 37°C for at most 6 passages in RPMI-1640 medium (Gibco, 11975093) containing 10% fetal bovine serum (FBS) (Life Technologies, 10082147) and 1% penicillin-streptomycin (Gibco, 15140122). $5 \times 10^5$ cells suspended in sterile Dulbecco's phosphate-buffered saline (DPBS) (Gibco, 14190144) were injected subcutaneously into both flanks of BALB/c mice (The Jackson Laboratory) between 7-8 weeks of age. Cells were tested free of mycoplasma and other pathogens by PCR panel testing (IMPACT II, IDEXX Bioanalytics, 41-00031)

Humanized PDX:

Humanized non-small cell lung cancer PDX tumors were derived by warm-passaging in which a PDX tumor was cut into 300 μm x 300 μm x 300 μm PDX LTFs and implanted in the hind flank of NSG mice (The Jackson Laboratory). When tumors reached ~50 $mm^3$, mice were injected intravenously with 10-12 x $10^6$ human peripheral blood mononuclear cells from a consenting healthy donor. Engraftment was confirmed by flow cytometry analysis of human CD45+ cells (>10%) in the peripheral blood.

Human tissue:

Protocols for the collection of human specimens were approved by an institutional review board. Resected tumors were collected by a waiver of consent or informed patient consent and core needle biopsies collected under informed patient consent. Tumor specimens were shipped overnight in NanoCool shipping containers (Peli Biothermal, 2-85225). The internal temperature of shipping systems was monitored using USB temperature data loggers. Figure S7 shows data from the shipment of 118 tumor specimens where >99% of specimens sustained a temperature between -2°C and 10°C for >98% of the shipping time. For studies including biomarker status, tumors were enrolled from patients with the following tumor types that have FDA-approved indications for αPD-1-treatment based on companion diagnostic biomarkers: colorectal, uterine, head and neck, and lung. At the end of the experiment, LTFs were fixed in formalin and histological assessment for PD-L1 or MMR protein expression was performed when PD-L1/MMR/MSI status was not available upon follow-up from patient charts.

**Specimen preparation**

Tumor Resections:

Resected tumors were attached to sample holders and embedded in a 3.8-4.0% w/v low-gelling temperature agarose (Sigma, A9045). The tumors were then carefully pushed through a cutting stage and cut into desired dimensions of resection LTFs (eg, length x width x thickness of 300 µm x 300 µm x 300 µm) by a fully automated, proprietary cutting device. Resection LTFs were collected directly into a 4°C bath containing cutting media (RPMI-1640 medium (Gibco, 11875093) supplemented with either 10% heat-inactivated human serum (Sigma, H3667) or 10% heat-inactivated FBS (Life Technologies 10082147), 10 mM HEPES (Gibco, 1630080), 1X MEM (Sigma Aldrich, M7145), 1% sodium pyruvate (Gibco, 11360070), 1% Penicillin-Streptomycin (Gibco, 15140122), and 1% GlutaMAX (Gibco, 35050061)). Resection LTFs were passed through a 420 µm, collected using a 200 µm filter and resuspended in a 50 mL conical tube with cutting media. Resection LTFs were next counted by an automated digital camera employing proprietary software. Resuspended fragments were gently pipetted to mix before aliquoting into a 24-well plate based on the target LTF number per well. To prevent T-cell egress (Voabil et al), 300 µL of a hydrogel (VitroGel®-3, TheWell Bioscience) was then added to each well and pipetted to mix with the LTFs. After the hydrogel polymerized, 1 mL of culture media containing a treatment was added to each well. The plate was imaged on the automated digital microscope to calculate the tissue volume present in each well. Resection LTFs were then maintained at 37°C and 5% $CO_2$ throughout experimentation.

Core Needle Biopsies:

12- to 20-gauge CNBs were measured for length, embedded in agarose solution, chilled on ice for 5 minutes to allow the agarose to solidify, and cut using an automated, proprietary cutting device into slices with a thickness of 300 µm. Biopsies used in the cross-well comparisons were cut cross-sectionally (90° angle to the longitudinal axis) and plated in a 8-well plate, whereas biopsies used in the sequential treatment studies were cut at the 20° angle to the longitudinal axis and plated in a 24-well culture plate. CNB LTFs were counted manually, equally distributed into the wells, and 300 µL of a hydrogel (VitroGel®-3, TheWell Bioscience; or a proprietary formula, Elephas) was added to each well. Fragments embedded in Elephas' proprietary hydrogel were exposed to a 395-nm UV light to polymerize and washed three times with DPBS. After hydrogel polymerization, 500 µL of culture media containing a treatment was added to each well with tissue and the plate was imaged on the automated digital camera to calculate the tissue volume present in each well. Specimens were maintained at 37°C and 5% $CO_2$ throughout experimentation.

**Treatments**

αCD3/αCD28 stimulation was performed with ImmunoCult™ Human CD3/CD28 T Cell Activator (STEMCELL Technologies Inc, 10971) at a final concentration of 25 µL/mL, except in experiments for Figure 5 when 100 µL/mL was used. This higher concentration was based on a pilot study which found the effect of αCD3/αCD28 stimulation reaches a plateau at this level (Data not shown). IgG and ICI antibodies were used at a concentration of 50 µg/mL and sourced from BioXCell. A control well was treated with a human IgG control antibody used at the final concentration of 50 µg/mL immediately after plating. The control for αCD3/αCD28 and αPD-1 was (RecombiMAb human IgG4 (S288P), CP147), αPD-L1 was a human IgG1 antibody (RecombiMAb human IgG1 (N297A), CP171) and αPD-1 and αCTLA-4 combination treatment was a combination of the human IgG4 antibody and a human IgG1 antibody (RecombiMAb human IgG1 isotype control, CP174). Exceptions to IgG matching include Colon 4834, Esophageal 4798 and Kidney 5023 control was IgG4 and Kidney 5327, IgG4+IgG1.

**Assays**

Viability and cytotoxicity assays:

The CCK8 reagent, WST-8 (water-soluble tetrazolium salt), was purchased from Abcam (228554). The reagent was diluted with culture medium and added to the culture well to obtain a final concentration of 0.1X at the start of the assay. At a specified time point, 60 µL of conditioned media containing the reagent was collected from a culture well into a clear, flat-bottom 96-well plate and absorbance was measured at 450 nm using a plate reader (BioLegend, 423555). Media containing CCK8 was used as a blank for the assay and relative viability was reported as the change in absorbance per hour.

LDH was measured using the LDH-Glo™ Cytotoxicity Assay kit (Promega, J2381). Conditioned media from a culture well was diluted 1:20 in the LDH storage buffer (200 mM Tris-HCl (ThermoFisher Scientific, J67501-AK), 10% Glycerol (Acros Organics, 327255000) and 1% BSA (Sigma Aldrich, A2058)) at the time of collection and frozen at -80°C for a minimum of 1 day before processing. Samples were thawed on ice and centrifuged before use. LDH Standard was diluted to the concentration of 32 mU/mL and serial dilutions were prepared with LDH storage buffer. Samples were added to a 96-well assay plate and mixed with LDH Detection Enzyme Mix and Reductase Substrate. The plate was incubated at room temperature for 30-60 minutes, protected from light, before being read for luminescence on a plate reader (PerkinElmer, EnSpire Alpha).

Optical Coherence Microscopy imaging:

A custom-designed optical coherence microscopy (OCM) system previously described (Liu et al 2024) and fitted with a 15X objective lens (Thorlabs, 0.70 NA), as well as accompanied software system (Elephas) was used to acquire time resolved, volumetric dynamic OCM (dOCM) data for viability assessment. The time series of each cross section was first registered to the center frame. For each pixel in the time series of volumetric OCM intensity I(x, y, z, t), fast Fourier transform was performed to obtain the power spectrum P(x, y, z, f). Intensity images were also acquired on the same system for structural analyses. Images were acquired every 24 hours for the same tumor specimen over 3 days of culture.

Cytokine profiling:

Using proprietary imaging software, the total fragment volume in each well was calculated to permit normalization of cytokine concentrations. Conditioned media collected from individual culture wells at defined time points were assessed using the Human XL Cytokine Luminex Performance Assay (R&D Systems, FCSTM18B, see supplemental table S4 for the list of analytes). The cytokine concentrations from samples were interpolated from a standard curve generated for each analyte. If the concentration was above the limit of quantitation, the value was set to the upper limit of quantitation; values below the limit of quantitation were unchanged and removed from analysis where noted. Additionally, six analytes (CD40L, EGF, FGF basic, IL-12p70, IL-15, and TNFβ) with concentrations consistently below LLOQ at 24 hours across samples were excluded from analysis. Replicate measurements were averaged unless otherwise noted. A cumulation of analyte concentrations was calculated to account for the portion of the analyte removed from the well at earlier time points whenever the earlier concentrations were assayed. For example, the analyte concentration at 4 hours was multiplied by the fractional supernatant draw relative to the total supernatant volume in the well and added to the concentration at 24 hours (the next time point) as shown in the equation below:

$$Cumulative\ Conc_{analyte} = Conc_{analyte}^{24hr} + Conc_{analyte}^{4hr} * \frac{100\ \mu L}{500\ \mu L}$$

Samples with multiple replicates in either the αPD-1 and IgG-treated group in the cross-well comparisons had their cumulative concentrations for each analyte averaged across all replicates. The sequential treatment configuration compared the cumulative concentrations from each replicate well separately.

Histology:

LTFs were fixed for at least 12 hours in 10% phosphate buffered formalin (Fisher Scientific SF100-4). Resection LTFs encapsulated in VitroGel®-3 were retrieved from the hydrogel after fixation and transferred to 70% alcohol before being pre-embedded in a 2% agarose solution (Sigma, A0576) in a custom-designed 3D printed mold. This procedure was designed to optimize the spatial arrangement of tumor fragments which produce a representative sampling of the tumor histology. In contrast, CNB LTFs encapsulated in Elephas hydrogel were directly transferred after fixation into 70% alcohol. Resection and CNB LTFs were paraffin-embedded in blocks and sectioned at 5-μm thickness on a rotary microtome (Leica, RM2255). Sections were mounted to slides using a tissue flotation water bath. Slides were then stained with hematoxylin and eosin (H&E) or processed for immunohistochemistry (IHC) and immunofluorescence (IF) using Ventana Medical Systems Discovery Ultra Autostainer (see table S5 for antibodies used). All quantification of histological slides was performed using Visiopharm software. Individual channels of mIF images were adjusted using Adobe Photoshop to reducing the effects of exposure-related intensity non-uniformities introduced during image acquisition and a global curve adjustment layer was applied to enhance overall image contrast. These adjustments were made solely to improve visual presentation, mitigate stitching-induced non-uniformities, and were not used for any quantitative analyses.

Statistical analyses:

Data was reported as mean ± standard deviation and statistical significance was assessed by Student's t-test or Analysis of Variance using Prism 10 software (GraphPad) unless otherwise noted. Fisher's exact test was performed in the Python SciPy software package [24].

For CNBs following the sequential treatment protocol, LTFs were treated with IgG at the start of culture and then with αPD-1 (lung specimens), αPD-L1 (gastrointestinal, GI specimens), or αPD-1 + αCTLA-4 (kidney specimens) at around 20 hours, such that LTFs in each well received control and ICI treatment sequentially. Conditioned media was collected at the ~4-, ~20- and ~48-hour time points. The rate of change (slope) of cumulative cytokine concentration (CC) was calculated for ICI and IgG treatment phases, as shown below.

$$\text{Slope}_{IgG} = \frac{CC_{20\ hour} - CC_{4\ hour}}{\Delta_{time}}$$

$$\text{Slope}_{ICI} = \frac{CC_{48\ hour} - CC_{20\ hour}}{\Delta_{time}}$$

The fold change of slope ($\text{Slope}_{FC}$) was then calculated, as shown below. If multiple replicate wells were tested for a given specimen, the maximum fold change of slope was reported.

$$\text{Slope}_{FC} = \frac{Slope_{ICI}}{Slope_{IgG}}$$

For resections, ICI-induced change in cytokine concentrations was determined by the difference in cumulative concentrations between ICI and IgG-treated wells (Delta), as shown below:

$$\Delta_{analyte} = CC_{analyte}^{\alpha PD1} - CC_{analyte}^{IgG}$$

To normalize the varying scale of Delta values across analytes and identify a potential responder to treatment, a trimmed-sample approach was applied. Calculations of the trimmed sample statistics occurred in two stages and leveraged the median (eg, $50^{th}$ percentile, E[X]) and median absolute difference (MAD) to describe the central tendency and spread of the population, respectively. The equation for MAD is shown below, where E denotes a median operator:

$$MAD = E[|X - E[X]|]$$

In the first stage, the median and MAD values were calculated using the full complement of samples. These initial values were used to identify inliers, with the definition of an inlier as any sample that falls within two MAD values of the median (ie, median - 2*MAD ≤ inlier ≤ median + 2*MAD). Using only the inliers, the median and MAD values were then recomputed to generate the trimmed statistics. The trimmed data was then transformed to Modified Z-scores (Mod. Z) [25] by the equation below:

$$Mod.Z = \frac{X - E[X]}{MAD} * 0.6745$$

This method allows for easier identification of upregulated cytokines through outliers relative to cytokines which exhibit no change or a decrease with αPD-1 treatment. The Modified Z transform differs from a traditional Z-score transform in that the central tendency and spread terms are derived from the median and MAD values, rather than the mean and standard deviation. The Mod. Z transform is known to be more robust to extreme-valued outliers due to the reliance on the median and MAD to define the transform [25], and is better optimized for outlier detection than a traditional Z-score transform.

Agglomerative hierarchical clustering was performed on the data, clustering both the individual samples (columns) and the analytes (rows) by Ward's method using TIBCO Spotfire™ software. The heatmap for changes in cytokine levels was generated using the Delta values for each analyte after statistical trimming and Mod. Z transformation of the cytokine dataset. Mod. Z-scores of -10 and 10 were selected as the lower and upper saturation points, respectively, as any value beyond this range already indicates meaningful analyte regulation. A conservative criterion to identify where upregulation occurred was defined by a Delta Mod. Z ≥ 5.0.
