## Supplemental figures and tables for "A live tumor fragment platform to assess immunotherapy response in core needle biopsies while addressing challenges of tumor heterogeneity"

#### Graphical Abstract

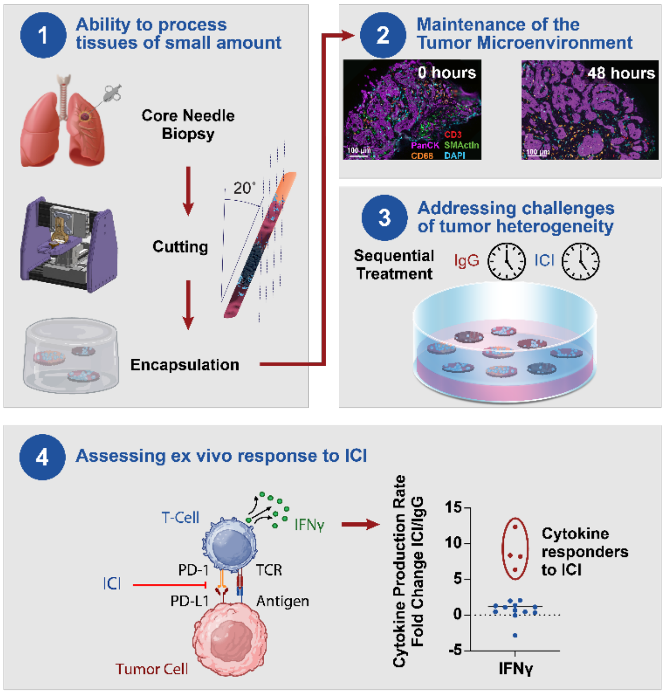


### A live tumor fragment platform to assess immunotherapy response in core needle biopsies while addressing challenges of tumor heterogeneity

**Authors**

T.S. Ramasubramanian, P. Adstamongkonkul, C.M. Scribano, C. Johnson, S. Caenepeel, L.C.F. Hrycyniak, L. Vedder, N. Dana, C. Baltes, T. Browning, Y. Chen, T. Dietz, E. Flietner, N. Kaplewski, A. Kellner, M. Korrer, C. Liu, N. Marhefke, P. McDonnell, A. Nasreen, T. Pope, A. Prasad, J. Richardson, S. Scheider, M. Schultz, C. Sood, A. Sunil, E. von Euw, E. Wait, E. Wargowski, P. Advani, B.J. Broome, A. Bruckbauer, A. Godwin, N. Kokabi, R.Martin, M. Robaina, G. Toia, J. Routh, A. Friedl, K. Eliceiri, M. Szulczewski, S. Johnson, J. Oliner, C. Capitini, D. Mukhopadhyay, J. Taube, D. Braun, H. J. Gierman

#### In brief

- An ex vivo platform for assessing ICI response using live tissue while preserving large areas of the contiguous native tumor microenviroment
- A sequential treatment strategy mitigates tumor heterogeneity challenges exacerbated when tissue amount is scarce
- The platform supports longitudinal profiling of tumor cytokine response

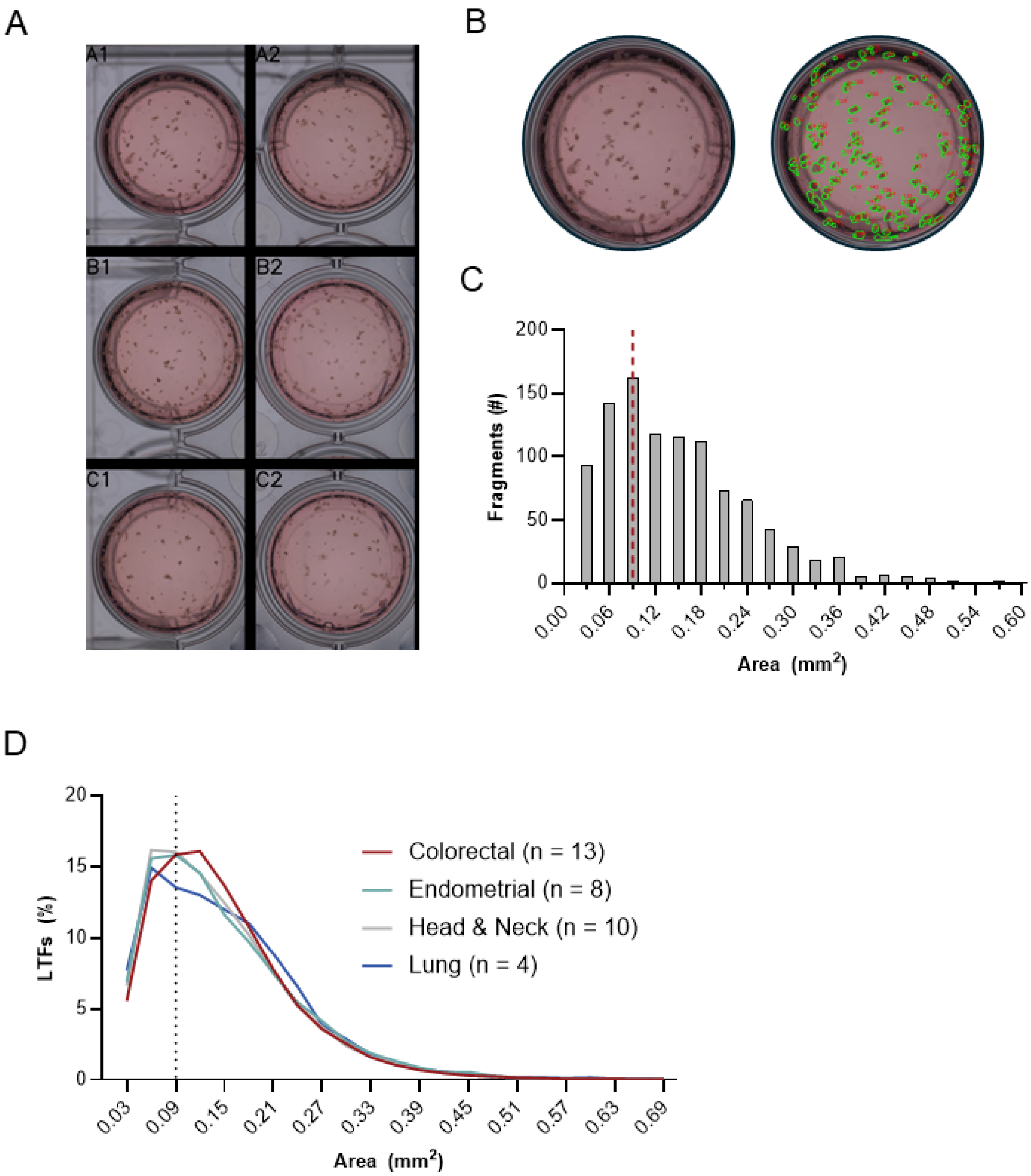
A
A1
A2
B1
B2
C1
C2
B
C
200
150
100
50
0
Fragments (#)
0.00
0.06
0.12
0.18
0.24
0.30
0.36
0.42
0.48
0.54
0.60
Area (mm$^2$)
D
20
15
10
5
0
LTFs (%)
Colorectal (n = 13)
Endometrial (n = 8)
Head & Neck (n = 10)
Lung (n = 4)
0.03
0.09
0.15
0.21
0.27
0.33
0.39
0.45
0.51
0.57
0.63
0.69
Area (mm$^2$)

**Figure S1:** Resected human tumors were cut using a proprietary cutting instrument which can cut tumor tissue automatically at user specified length, width and depth dimensions. Cutting tumors to 300 μm (length) x 300 μm (width) x 300 μm (depth) in size results in expected tissue areas across fragments which can then be randomly allocated to sample wells for treatment comparisons. **A** Resection LTFs were manually distributed amongst 6 wells of a culture plate after cutting. **B** An enlarged view of one of the wells (A1, left) and outlines of fragment area used for area calculations (right). **C** The number of fragments binned to a discrete range of areas are reported for the entire resected tumor. The dotted line represents the expected surface area for one face of a 300 μm x 300 μm x 300 μm cut fragment. **D** Histogram presenting the binned areas for fragments cut to 300 μm x 300 μm x 300 μm from colorectal (n=13), endometrial (n=8), head and neck (n=10) and lung (n=4) resected human tumors. Gray shading represents fragment areas within 1 standard deviation of the mean.

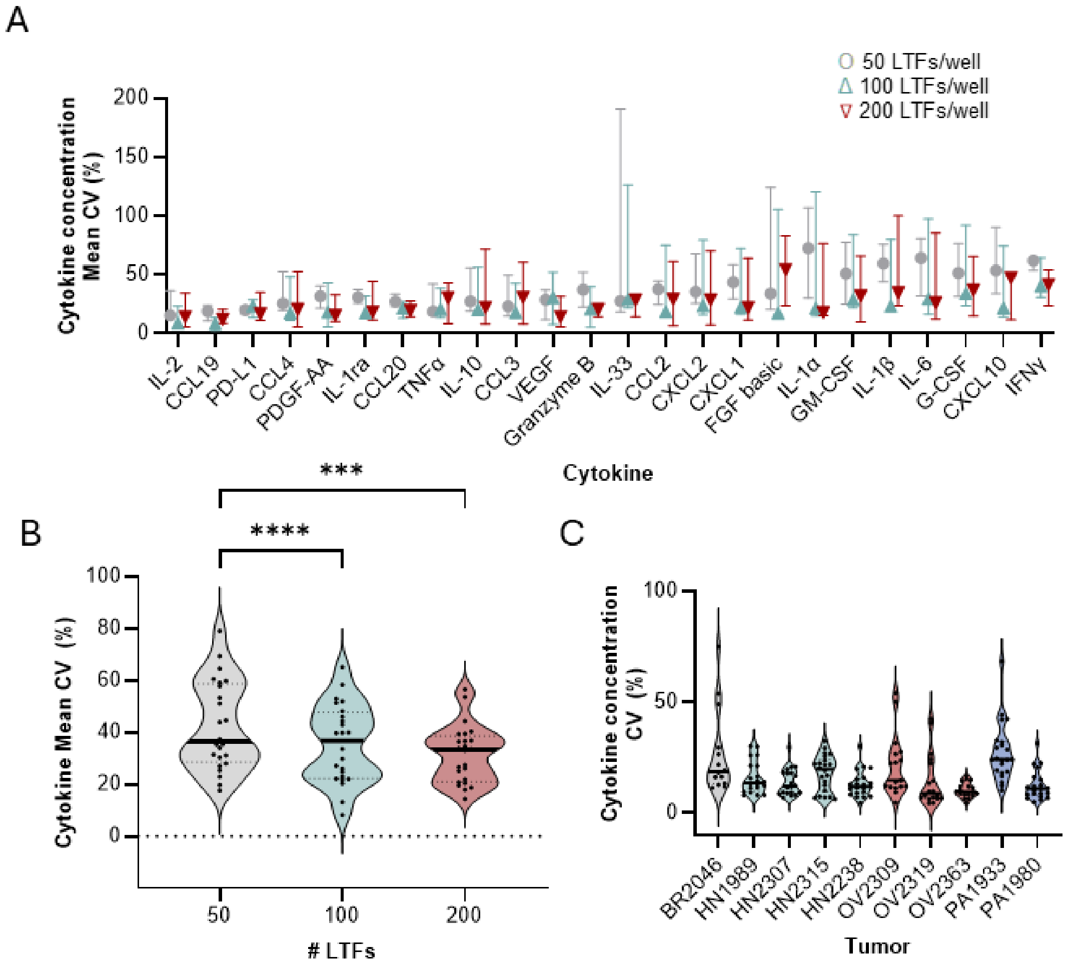
A
50 LTFs/well
100 LTFs/well
200 LTFs/well
Cytokine concentration Mean CV (%)
0
50
100
150
200
IL-2
CCL19
PD-L1
CCL4
PDGF-AA
IL-1ra
CCL20
TNFα
IL-10
CCL3
VEGF
Granzyme B
IL-33
CCL2
CXCL2
CXCL1
FGF basic
IL-1α
GM-CSF
IL-1β
IL-6
G-CSF
CXCL10
IFNγ
Cytokine
B
***
****
Cytokine Mean CV (%)
0
20
40
60
80
100
50
100
200
### LTFs
C
Cytokine concentration CV (%)
0
50
100
BR2046
HN1989
HN2307
HN2315
HN2238
OV2309
OV2319
OV2363
PA1933
PA1980
Tumor

**Figure S2**: **A** Effect of LTF number (~50, ~100 or ~200 LTFs/well) on variability (mean CV (%)) of cytokine induction (n=24 cytokines) following 24 hours of αCD3/αCD28 stimulation in 5 human tumor resections (lung, n=1; ovarian, n=1; colorectal, n=1; head and neck, n=2) each including 3 replicates reported per cytokine. **B** The effect of LTF number is further shown in a summary of CV (%) for cytokines across density of LTFs where a decrease in variance is shown between 50 LTFs and higher LTF densities. **C** Variability (CV (%)) across 5 replicate wells, each containing ~200 resection LTFs cut from 10 human tumors (pancreatic, n=2; head and neck, n=4; ovarian, n=3; triple negative breast cancer, n=1) stimulated with αCD3/αCD28 for 48 hours. *** $P < 0.001$ **** $P < 0.0001$

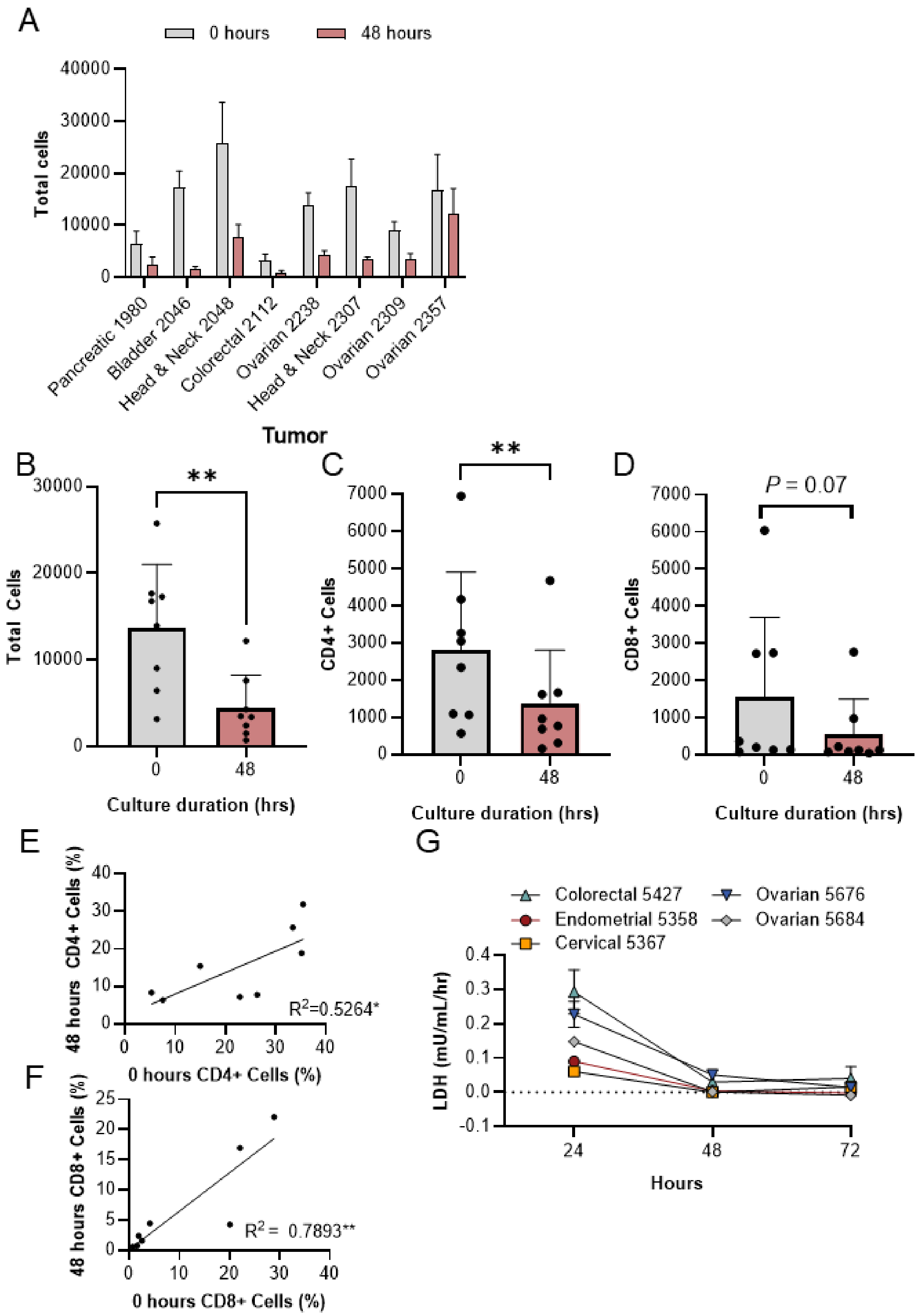
A
0 hours
48 hours
Total cells
40000
30000
20000
10000
0
Pancreatic 1980
Bladder 2046
Head & Neck 2048
Colorectal 2112
Ovarian 2238
Head & Neck 2307
Ovarian 2309
Ovarian 2357
Tumor
B
**
Total Cells
Culture duration (hrs)
C
CD4+ Cells
D
P = 0.07
CD8+ Cells
E
48 hours CD4+ Cells (%)
0 hours CD4+ Cells (%)
R2=0.5264*
F
48 hours CD8+ Cells (%)
0 hours CD8+ Cells (%)
R2 = 0.7893**
G
Colorectal 5427
Endometrial 5358
Cervical 5367
Ovarian 5676
Ovarian 5684
LDH (mU/mL/hr)
Hours

1

**Figure S3**: **A** Comparison of the total nuclei between 0 and 48 hours of culture**.** Summarized data for all specimens across culture duration for total cells (**B**), CD4+ (**C**), and CD8+ (**D**). The percentage of CD4-containing cells (**E)** and CD8-containing cells (**F**) show correlation between hours 0 and 48. **I** Changes in cytotoxicity (LDH assay) over 72 hours of ex vivo culture in LTFs derived from 5 human tumor resections shows a decrease in cytotoxicity between 24 and 48 hours, followed by stabilization between 48 and 72 hours of culture. * $P < 0.05$; ** $P < 0.01$.

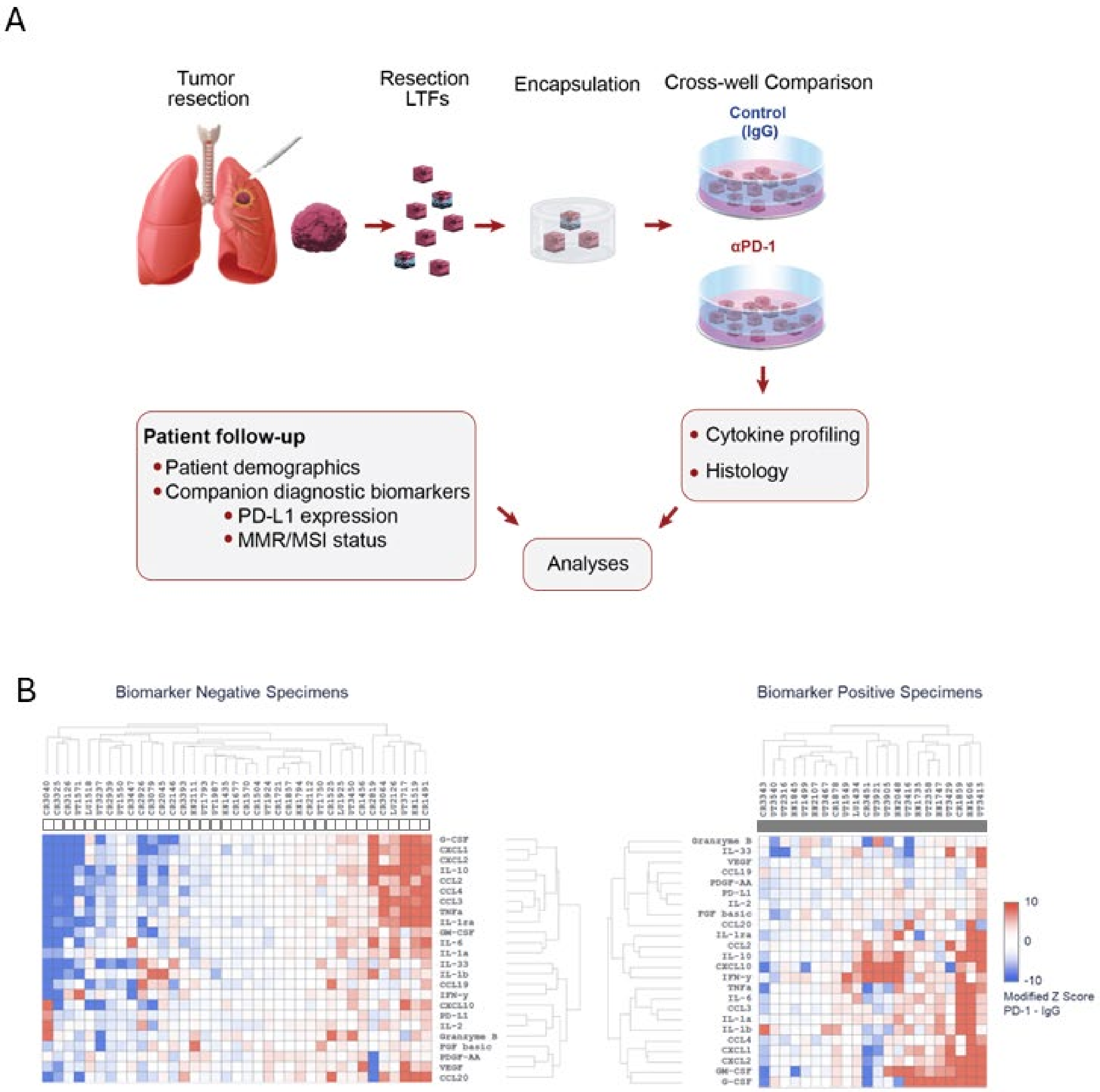
A
Tumor resection
Resection LTFs
Encapsulation
Cross-well Comparison
Control (IgG)
αPD-1
Cytokine profiling
Histology
Patient follow-up
Patient demographics
Companion diagnostic biomarkers
PD-L1 expression
MMR/MSI status
Analyses
B
Biomarker Negative Specimens
Biomarker Positive Specimens
G-CSF
CXCL1
CXCL2
IL-10
CCL2
CCL4
CCL3
TNFa
IL-1ra
GM-CSF
IL-6
IL-1a
IL-33
IL-1b
CCL19
IFN-y
CXCL10
PD-L1
IL-2
Granzyme B
FGF basic
PDGF-AA
VEGF
CCL20
10
0
-10
Modified Z Score
PD-1 - IgG

**Figure S4 A** Schematic showing the methods used for creating resection LTFs to assess cytokine response from PD-L1/MMR/MSI positive (PD-L1+/dMMR/MSI-high) and PD-L1/MMR/MSI negative (PD-L1-/pMMR/MSS) specimens. **B** Unsupervised hierarchical clustering of cytokine profiles from patient tumor resections using modified Z-scores of the difference in cytokine concentrations between the ICI and IgG treated groups where PD-L1/MMR/MSI-positive and PD-L1/MMR/MSI-negative specimens are presented separately.

A

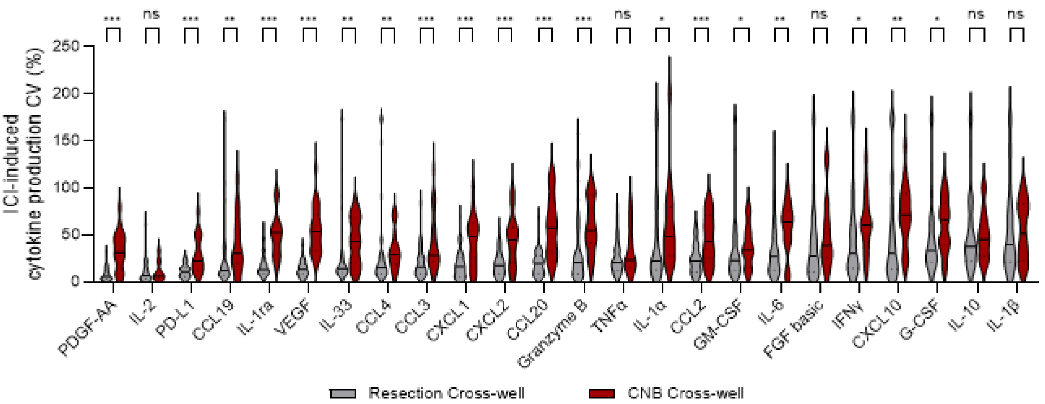

ICI-induced cytokine production CV (%)
0
50
100
150
200
250
*** ns *** ** *** *** ** ** *** *** *** *** *** ns * *** * ** ns * ** * ns ns
PDGF-AA
IL-2
PD-L1
CCL19
IL-1ra
VEGF
IL-33
CCL4
CCL3
CXCL1
CXCL2
CCL20
Granzyme B
TNFα
IL-1α
CCL2
GM-CSF
IL-6
FGF basic
IFNγ
CXCL10
G-CSF
IL-10
IL-1β
Resection Cross-well
CNB Cross-well

**Figure S5:** Variance (CV (%) of ICI-induced cytokine production in resection and CNB cross well experiments represented for each cytokine measured. * $P < 0.05$, ** $P < 0.005$, *** $P < 0.001$

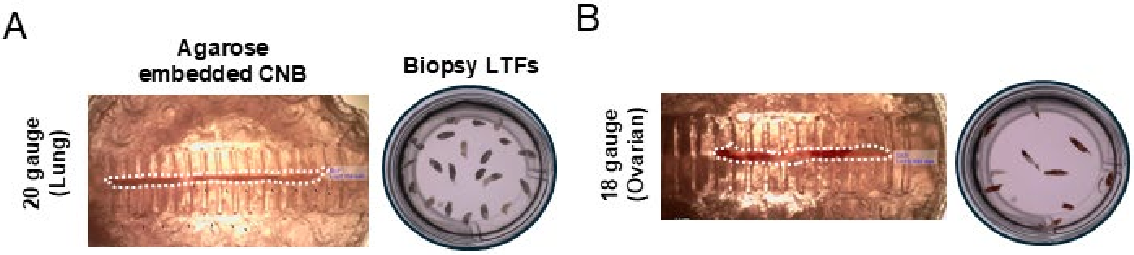
A
Agarose embedded CNB
Biopsy LTFs
20 gauge (Lung)
B
18 gauge (Ovarian)

**Figure S6:** Representative images showing a lung tumor CNB specimen from a 20-gauge biopsy needle (**A**) and an ovarian tumor CNB specimen from an 18-gauge biopsy needle (**B**) embedded in agarose in preparation for cutting (left) and resulting CNB LTFs post cutting (right).

A

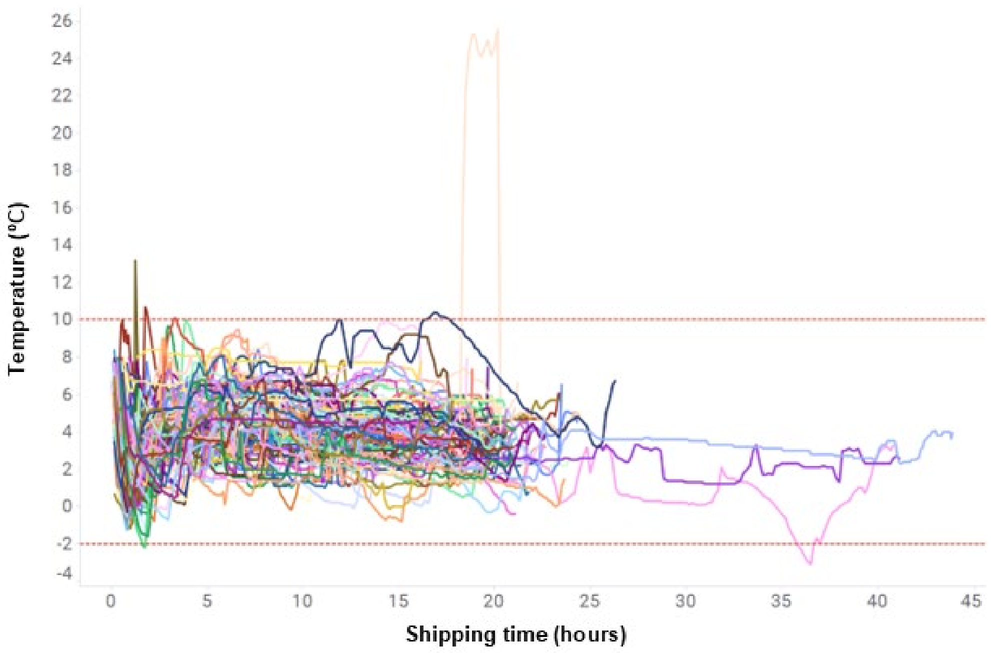
Temperature (ºC)
26
24
22
20
18
16
14
12
10
8
6
4
2
0
-2
-4
0
5
10
15
20
25
30
35
40
45
Shipping time (hours)

**Figure S7:** Internal temperature of NanoCool shipping system vs. shipping time for 118 tumor specimens collected consecutively shows tight temperature regulation during overnight shipping of CNBs from diverse clinical sites.

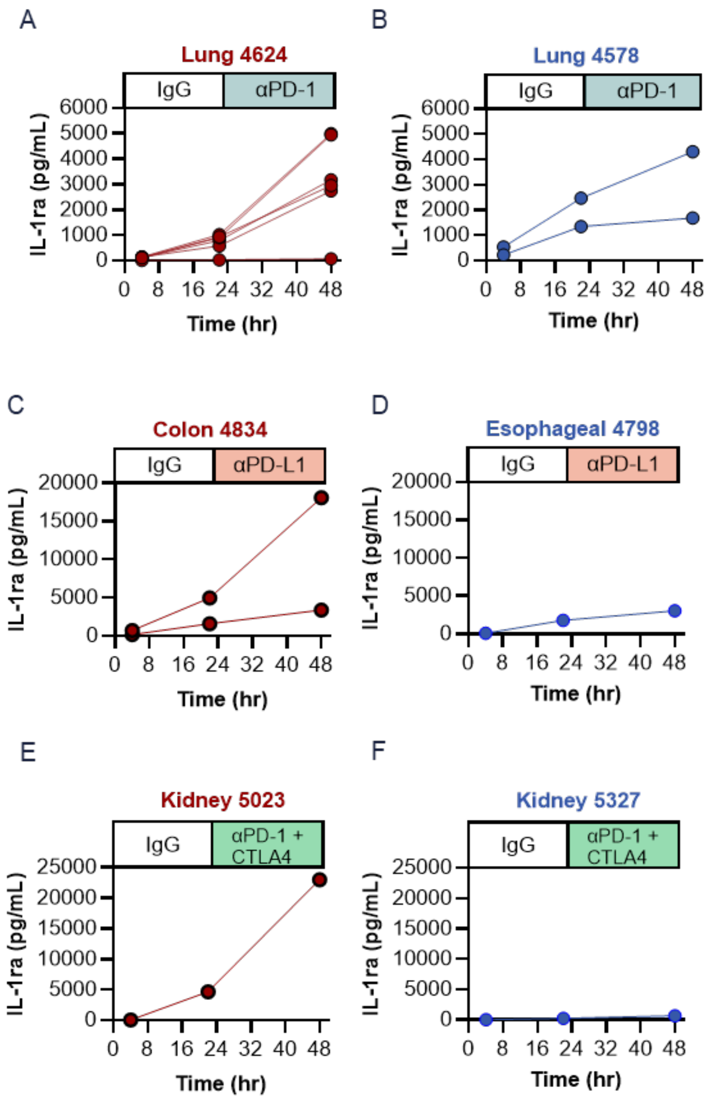
A
Lung 4624
IgG
αPD-1
IL-1ra (pg/mL)
Time (hr)
B
Lung 4578
IgG
αPD-1
IL-1ra (pg/mL)
Time (hr)
C
Colon 4834
IgG
αPD-L1
IL-1ra (pg/mL)
Time (hr)
D
Esophageal 4798
IgG
αPD-L1
IL-1ra (pg/mL)
Time (hr)
E
Kidney 5023
IgG
αPD-1 + CTLA4
IL-1ra (pg/mL)
Time (hr)
F
Kidney 5327
IgG
αPD-1 + CTLA4
IL-1ra (pg/mL)
Time (hr)

**Figure S8 A** CCK8 analysis of CNB LTFs derived from 4 human tumor resections demonstrates maintenance of viability over 72 hours of culture. **B** Cytotoxicity evaluation (LDH assay) over 72 hours of ex vivo culture in LTFs derived from 4 human tumor CNBs shows a decrease in cytotoxicity between 24 and 48 hours, followed by stabilization between 48 and 72 hours of culture, similar to what was shown in resection LTFs. **C** H&E-stained sections of a human liver cancer (Liver 5605) biopsy LTF encapsulated in Elephas hydrogel and cultured for 0 (upper) or 48 (lower) hours. Both 0-hour and 48-hour CNB LTFs show islands of carcinoma cells (green arrow) within sparse stroma (matrix-enriched stroma, orange arrow; collagen-rich stroma, yellow arrow), lymphocytes (red arrows) and fibroblasts (blue arrow).

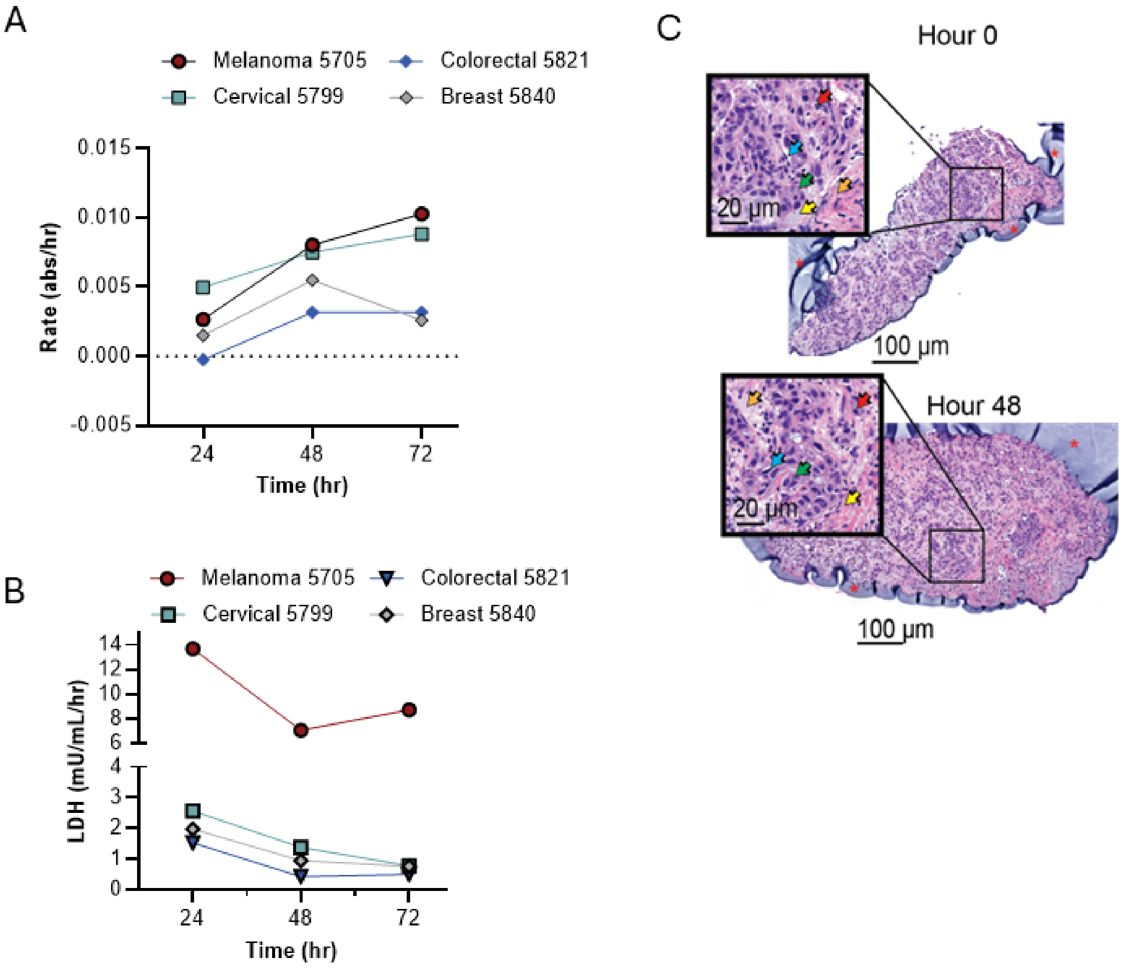
A
Melanoma 5705
Colorectal 5821
Cervical 5799
Breast 5840
Rate (abs/hr)
0.015
0.010
0.005
0.000
-0.005
24
48
72
Time (hr)
B
Melanoma 5705
Colorectal 5821
Cervical 5799
Breast 5840
LDH (mU/mL/hr)
14
12
10
8
6
4
3
2
1
0
24
48
72
Time (hr)
C
Hour 0
20 μm
100 μm
Hour 48
20 μm
100 μm

**Figure S9** IL-1Ra represents a cytokine for which little to no change in cytokine production rate was observed between the IgG and ICI treatment phases for all specimens profiled. IL-1Ra measured at 4, 24 and 48 hours for specimens which showed an increase in IFNγ and CXCL10 induction following ICI treatment (αPD-1, **A**; αPD-L1,**C;** αPD-1+ αCTLA-4, **E**) and specimens which showed no change in IFNγ and CXCL10 induction following ICI treatment (αPD-1, **B**; αPD-L1, **D;** αPD-1+ αCTLA-4, **F**). IL-1Ra shows robust concentrations with a smaller induction rate during the IgG and ICI treatment phases.

**SUPPLEMENTAL TABLES**

**Table S1:** Table of patient characteristics showing patient demographics, diagnoses, companion diagnostic biomarker status, clinical stage and treatments for biomarker resection LTF study.

| Patient Characteristics | | | |
|---|---|---|---|
| | | N | % |
| Sex | M | 21 | 35.6% |
| | F | 30 | 50.8% |
| | Unknown | 8 | 13.6% |
| Age | Range | 37-88 years | |
| | 30-40 | 2 | 3.4% |
| | 41-50 | 9 | 15.3% |
| | 51-60 | 8 | 13.6% |
| | 61-70 | 15 | 25.4% |
| | 71-80 | 9 | 15.3% |
| | 81+ | 7 | 11.9% |
| | Unknown | 9 | 15.3% |
| Race | White | 41 | 68.3% |
| | Black | 6 | 10.2% |
| | American Indian or Alaska Native | 1 | 1.7% |
| | Unknown | 11 | 18.6% |
| Tumor Type | Lung | 10 | 16.9% |
| | Head & Neck | 4 | 6.8% |
| | Colorectal | 25 | 42.4% |
| | Uterine | 20 | 33.9% |
| | Primary | 42 | 71.2% |
| | Secondary | 17 | 28.8% |
| Companion diagnostic biomarker status | Positive | 22 | 37.3% |
| | PD-L1 | 7 | 31.8% |
| | MSI | 15 | 68.2% |
| | Negative | 37 | 62.7% |
| | PD-L1 | 7 | 18.9% |
| | MSI | 30 | 81.1 |
| Clinical Stage | 1 | 5 | 8.5% |
| | 2 | 6 | 10.2% |
| | 3 | 9 | 15.3% |
| | 4 | 6 | 10.2% |
| | Unknown | 33 | 55.9% |

| IO Treatment | | | |
|---|---|---|---|
| Neoadjuvant IO | Yes | 2 | 3.4% |
| | No | 20 | 33.9% |
| | Unknown | 37 | 62.7% |
| Adjuvant IO | Yes | 4 | 6.8% |
| | No | 35 | 59.3% |
| | Unknown | 20 | 33.9% |
| Chemotherapy | Yes | 7 | 11.9% |
| | No | 21 | 35.6% |
| | Unknown | 31 | 52.5% |

**Table S2:** Table of cytokines showing increased rate of induction in ICI treated compared to IgG treated LTFs from PD-L1/MMR/MSI positive and PD-L1/MMR/MSI negative specimens with statistical comparisons. GM-CSF, IFNγ and CXCL10 all show significantly increased rates of cytokine production in the PD-L1/MMR/MSI positive compared to PD-L1/MMR/MSI negative specimens.

| **Analyte** | **PD-L1/MMR/MSI positive rate** | **PD-L1/MMR/MSI negative rate** | ***P*-value** |
|---|---|---|---|
| GM-CSF*** | 0.41 | 0.03 | 0.0003 |
| IFNy ** | 0.36 | 0.03 | 0.001 |
| CXCL10* | 0.32 | 0.08 | 0.03 |
| G-CSF | 0.36 | 0.14 | 0.055 |
| IL-1b | 0.32 | 0.11 | 0.081 |
| CCL2 | 0.18 | 0.08 | 0.407 |
| CXCL2 | 0.18 | 0.11 | 0.455 |
| IL-1a | 0.18 | 0.11 | 0.455 |
| TNFa | 0.23 | 0.14 | 0.477 |
| CCL19 | 0.00 | 0.05 | 0.524 |
| IL-2 | 0.00 | 0.05 | 0.524 |
| IL-33 | 0.09 | 0.05 | 0.624 |
| CCL20 | 0.09 | 0.16 | 0.697 |
| CCL3 | 0.09 | 0.14 | 0.702 |
| CXCL1 | 0.18 | 0.14 | 0.715 |
| IL-10 | 0.23 | 0.16 | 0.731 |
| CCL4 | 0.14 | 0.11 | 1 |
| FGF basic | 0.05 | 0.05 | 1 |
| Granzyme B | 0.05 | 0.08 | 1 |
| IL-1ra | 0.14 | 0.14 | 1 |
| IL-6 | 0.18 | 0.16 | 1 |
| PD-L1 | 0.00 | 0.03 | 1 |
| PDGF-AA | 0.00 | 0.00 | 1 |
| VEGF | 0.05 | 0.05 | 1 |

**Table S3:** Estimates for the target total tissue per well for resection LTFs cut to 300 µm cubes, CNB LTFs cut at a 90º or 20º angle from samples collected by various needle gauges.

| LTF characteristics | | | | | | | | | | | |
|---|---|---|---|---|---|---|---|---|---|---|---|
| | | Shape | L (mm) | W (mm) | H (mm) | Diameter (mm) | Long axis (mm) | Short axis (mm) | Volume per fragment ($mm^3$) | # frags/well | Total tissue volume ($mm^3$) |
| **Resections** | | | | | | | | | | | |
| Scored and cut | | Cuboid | 0.3 | 0.3 | 0.3 | | | | 0.027 | 200 | 5.400 |
| **Biopsies** | | | | | | | | | | | |
| 90° cut | 12 g | Circular | | | 0.3 | 2.2 | | | 1.140 | 5 | 5.702 |
| | 14 g | Circular | | | 0.3 | 1.6 | | | 0.603 | 9 | 5.429 |
| | 16 g | Circular | | | 0.3 | 1.2 | | | 0.339 | 16 | 5.429 |
| | 18 g | Circular | | | 0.3 | 0.84 | | | 0.166 | 33 | 5.486 |
| | 20 g | Circular | | | 0.3 | 0.6 | | | 0.085 | 64 | 5.429 |
| 20° cut | 12 g | Oval | | | 0.3 | | 6.11 | 2.07 | 2.980 | 2 | 5.960 |
| | 14 g | Oval | | | 0.3 | | 4.44 | 1.51 | 1.580 | 3 | 4.739 |
| | 16 g | Oval | | | 0.3 | | 3.33 | 1.13 | 0.887 | 6 | 5.320 |
| | 18 g | Oval | | | 0.3 | | 2.33 | 0.8 | 0.439 | 12 | 5.270 |
| | 20 g | Oval | | | 0.3 | | 1.66 | 0.56 | 0.219 | 25 | 5.47 |

**Table S4:** List of cytokines assayed. Cytokines denoted with an asterisk (*) were often not detected and were therefore excluded from analysis.

| Analytes in the 30 plex Luminex assay | |
|---|---|
| CCL19/MIP-3β | Lymphotoxin-α/TNF-β* |
| CCL2/JE/MCP-1 | IL-10 |
| CCL20/MIP-3α | IL-12 p70* |
| PD-L1/B7-H1 | IL-15* |
| CCL3/MIP-1α | IL-1β/IL-1F2 |
| CCL4/MIP-1β | IL-1ra/IL-1F3 |
| CD40 Ligand/TNFSF5* | IL-2 |
| CXCL1/GROα/KC/CINC-1 | IL-33 |
| CXCL10/IP-10/CRG-2 | IL-6 |
| CXCL2/GROβ/MIP-2/CINC-3 | PDGF-AA |
| EGF* | TNF-α |
| IFN-γ | VEGF |
| Flt-3 Ligand/FLT3L | FGF basic/FGF2/bFGF* |
| G-CSF | IL-1α/IL-1F1 |
| GM-CSF | Granzyme B |

**Table S5:** List of antibodies used for IHC and immunofluorescence experiments.

| Antibody | Clone | Manufacturer and Catalog Number |
|---|---|---|
| Actin, smooth muscle | 1A4 | Cell Marque #202M |
| anti-CD3 | 2GV6 | Ventana Medical Systems #790-4341 |
| anti-CD4 | SP35 | Ventana Medical Systems #790-4423 |
| anti-CD8 | SP57 | Ventana Medical Systems #790-4460 |
| anti-CD68 | KP-1 | Ventana Medical Systems #790-2931 |
| anti-Keratin, pan | AE1/AE3& PCK26 | Ventana Medical Systems #760-2595 |
